# Canalization of flower production across thermal environments requires Florigen and CLAVATA signaling

**DOI:** 10.1101/2025.03.23.644808

**Authors:** Elizabeth S. Smith, Amala John, Andrew C. Willoughby, Daniel S. Jones, Vinicius C. Galvão, Christian Fankhauser, Zachary L. Nimchuk

## Abstract

The ability to maintain invariant developmental phenotypes across disparate environments is termed canalization, but few examples of canalization mechanisms are described. In plants, robust flower production across environmental gradients contributes to reproductive success and agricultural yields. Flowers are produced by the shoot apical meristem (SAM) in an auxin-dependent manner following the switch from vegetative growth to the reproductive phase. While the timing of this phase change, called the floral transition, is sensitized to numerous environmental and endogenous signals, flower formation itself is remarkably invariant across environmental conditions. Previously we found that CLAVATA peptide signaling promotes auxin-dependent flower primordia formation in cool environments, but that high temperatures can restore primordia formation through unknown mechanisms. Here, we show that heat promotes floral primordia patterning and formation in SAMs not by increased auxin production, but through the production of the mobile flowering signal, florigen, in leaves. Florigen, which includes *FLOWERING LOCUS T* (*FT*) and its paralog *TWIN SISTER OF FT* (*TSF*) in *Arabidopsis thaliana*, is necessary and sufficient to buffer flower production against the loss of CLAVATA signaling and promotes heat-mediated primordia formation through specific SAM expressed transcriptional regulators. We find that sustained florigen production is necessary for continuous flower primordia production at warmer temperatures, contrasting florigen’s switch-like control of floral transition. Lastly, we show that CLAVATA signaling and florigen synergize to canalize flower production across broad temperature ranges. This work sheds light on the mechanisms governing the canalization of plant development and provides potential targets for engineering crop plants with improved thermal tolerances.

## Introduction

Many developmental traits are buffered against disruption by genetic and environmental perturbations. This phenomenon, originally termed “canalization” by Waddington, provides a selective advantage by ensuring that fluctuations in environmental conditions and stochastic mutations do not interfere with developmental processes essential for reproduction^1,2^. In plants, successful reproduction depends on robust flower production. The timing of the switch from vegetative growth to the reproductive phase, termed the floral transition, varies widely within a species in response to local environmental conditions^3^. In contrast, the production of normal flowers following the floral transition is remarkably invariant. This suggests that the pathways governing flower formation are highly canalized. As many agricultural yields are inherently linked to flower production, and temperature fluctuations can negatively impact crop yields, understanding the pathways that canalize flower production may enable engineering of crop species to optimize flower production and yield^4,5^.

Moderately elevated temperatures induce diverse developmental and growth changes in plants termed thermomorphogenesis. In *Arabidopsis thaliana,* thermomorphogenesis in above ground tissues includes increases in hypocotyl, leaf petiole, and stem elongation, and leaf hyponasty^6^. In particular, thermomorphogenic growth in hypocotyls and leaves is facilitated by increased auxin biosynthesis at elevated temperatures (ca. 27-30°C)^7^. Curiously, flower primordia production in SAM tissue proceeds normally across temperature ranges typically assayed during thermomorphogenesis experiments, despite flower primordia formation being critically dependent on auxin^8–11^. How heat impacts SAM functions, and whether thermomorphogenesis circuits operate similarly in SAMs compared to other tissues is an unresolved question.

Recent work from our lab showed that auxin-dependent primordia outgrowth at cooler temperatures requires CLAVATA3/EMBRYO SURROUNDING REGION peptide (CLEp) signaling^12,13^. This function is genetically separable from the well-described role of CLEp signaling in restricting stem cell proliferation in the SAM by buffering expression of *WUSCHEL* (*WUS*), a transcription factor promoting stem cell maintenance^14,15^. This work revealed that CLAVATA3 peptide (CLV3p) promotes primordia outgrowth by signaling through the CLEp receptors CLAVATA1 (CLV1) and the obligate dimer of CORYNE (CRN) and CLAVATA2 (CLV2), which encode a transmembrane pseudokinase and an extracellular leucine-rich repeat (LRR) receptor-like protein, respectively. In cool temperatures, *crn*/*clv2* plants bolt and produce 2-5 normal flowers before entering a prolonged period in which floral primordia form, but primordia outgrowth ceases at floral stage 3 (before floral meristem formation) and shoot elongation stalls (Figure S1A). After the production of approximately 30 terminated primordia during this “termination phase”, normal flower formation and shoot elongation resume for unknown reasons. Notably, high temperatures restore normal patterns of auxin-dependent floral primordia formation and outgrowth, and thus flower production, to *crn*/*clv2* plants^12,13^. How elevated temperatures buffer flower formation is unclear, and whether this involves similar thermomorphogenic circuitry as in other plant tissues remains unknown. Here, we show that canalized flower production is facilitated by elevated expression levels of the mobile signal florigen induced during thermomorphogenesis, and not by auxin-mediated pathways as in other tissues. We show that elevated expression levels of *FT*, a component of florigen, are necessary and sufficient to bypass the requirement for CLEp signaling during flower production at both cool and hot temperatures. This reveals that sustained *FT* expression is essential during floral primordia initiation and outgrowth, distinct from its well-described role in triggering floral transition. Lastly, we reveal that continuous flower production requires simultaneous florigen and CLEp signaling across temperature regimes. Collectively, these data reveal that dual developmental signaling programs promote the canalization of reproductive development against environmental perturbations, raising the possibility these pathways could be engineered to buffer crop development across temperature ranges.

## Results

### The transcriptional circuitry required for heat-induced auxin biosynthesis is dispensable for canalized flower formation at elevated temperatures

Previously we observed that aborted floral primordia in cool-grown *crn* SAMs showed reduced auxin-induced gene expression, consistent with the essential role of auxin in promoting primordia specification and outgrowth^11,13^. Here we focus on *crn*/*clv2* plants, as stem cell over-proliferation in other *clv* mutants obscures floral primordia phenotypes.^12^ As thermomorphogenic growth in other tissues results from increased auxin biosynthesis at high temperatures, we initially suspected that elevated levels of auxin induced during thermomorphogenesis canalized flower production in *crn* mutants at high temperatures^7^. Increased auxin biosynthesis at high temperatures is regulated by the thermosensing transcription factor EARLY FLOWERING3 (ELF3). In cooler temperatures, ELF3 transcriptionally represses *PHYTOCHOROME INTERACTING FACTOR 4* (*PIF4*), a transcription factor that promotes the expression of *YUCCA8* (*YUC8*), a member of the YUC family of rate-limiting auxin biosynthesis enzymes^16,17^. ELF3 forms condensates at high temperatures, permitting increased *PIF4* and *YUC8* expression, leading to auxin accumulation (Figure S1B)^18^. Based on our previous observation that *elf3* null mutations, which mimic warmer growth conditions, restore flower formation to *crn* plants at cooler temperatures, we suspected that ELF3-mediated rewiring of auxin biosynthesis facilitated heat-induced flower formation in *crn*^13^. To test this, we asked whether *PIF4* or *YUC8* were necessary for the restoration of floral primordia outgrowth in *crn elf3* double mutants grown in cooler temperatures (17-18°C). To quantify flower formation, we classified the first thirty flower attempts as normal (all floral organs present), terminated flower (pedicel forms but gynoecium is missing), or terminated primordia (no pedicel or floral organ formation), as previously described^12,13^. While *crn elf3 pif4-2* plants displayed modestly increased rates of primordia termination when compared to *crn elf3,* neither *crn elf3 yuc8-1* nor *crn elf3 pif4-2* mutant combinations displayed terminated primordia numbers similar to *crn* plants, suggesting that *elf3*-mediated restoration of primordia formation in *crn* plants is largely independent of *YUC8,* with only minor contributions from *PIF4* (Figures S1C-D).

As additional *PIF* and *YUC* paralogs can act redundantly during thermomorphogenesis, we asked whether heat could restore flower formation to *crn-10A pif34578* and *crn-10A yuc289* higher order mutants ^19–21^ (see materials and methods). At cool temperatures, *pif34578* did not affect primordia termination in *crn-10A,* while *crn-10A yuc289* showed a minor increase in flower formation, indicating that neither of these sets of *PIF* or Y*UC* paralogs play a major role in primordia outgrowth at cool temperatures (Figures S1E-F). Surprisingly, heat completely restored floral primordia formation to *crn-10A pif34589* and *crn-10A yuc289* mutants, indicating that the transcriptional circuitry required for increased auxin accumulation downstream of ELF3 thermosensing in other tissues is dispensable for heat-induced flower formation in *crn* plants (Figures S1G-H). This suggests that heat acts via alternative mechanisms during the canalization of flower production in the SAM.

### Acceleration of the floral transition restores primordia formation to *crn* mutants

As the ELF3-dependent transcriptional circuit that promotes auxin biosynthesis during thermomorphogenesis is dispensable for heat-mediated canalization of flower production, we sought to identify whether other transcriptional networks facilitate heat-mediated primordia formation. We noticed that the heat-labile MADS Box transcription factor SHORT VEGETATIVE PHASE (SVP), which represses floral transition in response to fluctuations in environmental temperature, is upregulated in *crn* SAMs during the primordia termination phase in previous RNA-Seq analyses (Figure 1A)^22,23^. Mirroring this, we found modestly increased SVP accumulation in cool-grown *crn* SAMs using a functional *pSVP::SVP-GFP* reporter^24^. While ectopic SVP levels were low in *crn* SAM cells, SVP displayed apparent nuclear localization, while SVP was nearly undetectable in Col-0 SAMs proper (Figure1B). We therefore suspected that elevated SVP in *crn* may be associated with primordia termination, and that SVP degradation in high temperatures may be linked to the restoration of normal flower formation. To first test this, we genetically mimicked high temperatures by mutating *svp* in the *crn* background. Indeed, *crn svp* double mutants grown in cooler temperatures displayed normal flower production, supporting a role for SVP accumulation in blocking primordia outgrowth in *crn* plants, and suggesting that SVP degradation in heat may explain the restoration of floral primordia outgrowth (Figures 1C-E).

**Figure 1.**
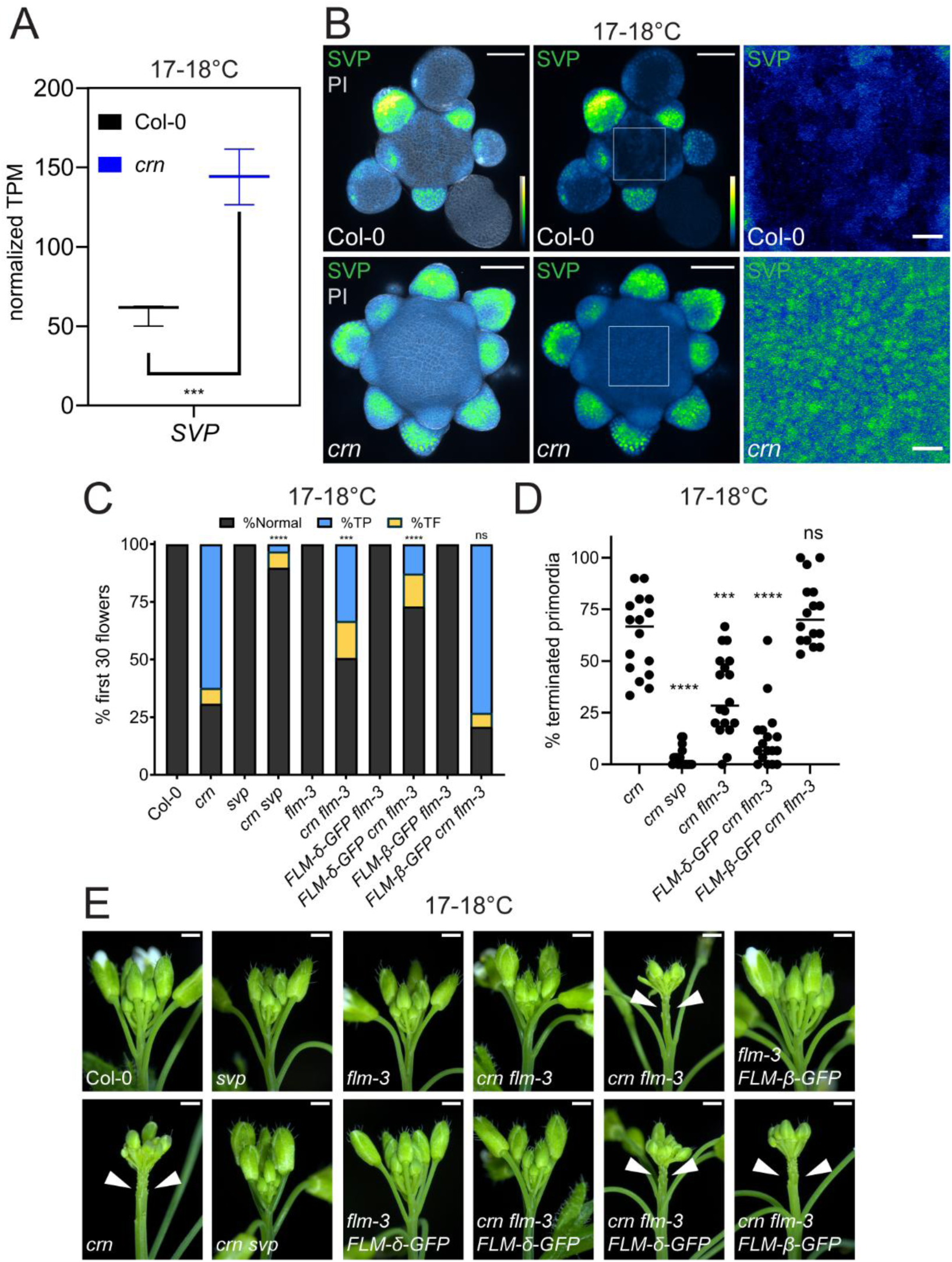
Loss of SVP restores primordia outgrowth to *crn* at cool temperatures. (A) Expression levels of *SVP* from RNA-Seq analysis in Col-0 and *crn* SAMs grown at 17-18°C. Normalized transcripts per million (TPM) summed for all three *SVP* isoforms for three biological replicates are plotted from Kallisto/Sleuth RNA-Seq analysis. The line indicates the median. P values for the three isoforms of *SVP* were p=0.000852, p=0.000266, and p<0.0001. (B) Maximum intensity projection of *pSVP::SVP-GFP* (GFP signal is color-coded for intensity using the BlueToYellow LUT) with PI (Gray) and *pSVP::SVP-GFP* alone in Col-0 (n=8) and *crn* (n=7) SAMs grown at 17-18°C. Square on SAMs in middle column indicates zoomed in region shown in right column. Scale bars are 50 µM (left and middle columns) and 10 µM (right column). Brightness and contrast were adjusted identically for both images in the right column compared to the left and middle columns to illustrate difference in expression levels between Col-0 and *crn*. (C) Quantification of terminated floral primordia in Col-0 (n=17), *crn* (n=16), *svp* (n=16), *crn svp* (n=18), *flm-3* (n=18), *crn flm-3* (n=18), *FLM-δ-GFP flm-3* (n=17), *FLM-δ-GFP crn flm-3* (n=17), *FLM-β-GFP flm-3* (n=15), and *FLM-β-GFP crn flm-3* (n=16) grown at 17-18°C. Statistical comparisons to *crn* are indicated on the graph. (D) %Terminated primordia for each of the *crn, crn svp, crn flm-3, FLM-δ-GFP crn flm-3* and *FLM-β-GFP crn flm-3* individuals summarized in (D). Statistical comparisons to *crn* are indicated on the graph. Line indicates the median. Statistical significance between %terminated primordia in indicated pairwise comparisons was calculated with multiple Mann-Whitney tests using the Holm-Šídák method for correcting multiple comparisons. Significance is represented on the graph using asterisks (p < 0.05 *; p < 0.01 **; p < 0.001 ***; p < 0.0001****.) (E) Representative inflorescence images of genotypes quantified in (C) and (D). Scale bars are 1 mm. Arrowheads indicate terminated floral primordia.

To further explore this possibility, we took advantage of the MADS Box transcription factor FLOWERING LOCUS M (FLM), which directly mediates the effects of temperature on SVP stability^23,25,26^. At cooler temperatures, the *FLM-β* isoform is predominately expressed and facilitates nuclear accumulation of SVP and direct repression of the transcriptional programs that induce floral transition. At higher temperatures, SVP is targeted for proteasomal degradation by the more abundant FLM-δ isoform, thereby permitting floral transition^23,25,26^. To test if the regulation of SVP accumulation by FLM influences primordia formation, we first asked whether the expression of specific *FLM* isoforms impacts flower formation in *crn* plants at cooler temperatures, using previously published *FLM-GFP* isoforms expressed from the native *FLM* promoter^23^. Indeed, *crn flm-3* expressing *FLM-δ-GFP* displayed almost normal flower formation at cooler temperatures, similar to *crn svp* double mutants and consistent with the role of FLM-δ in promoting SVP degradation. In contrast, the null *flm-3* partially restored flower formation to *crn* while *FLM-β-GFP crn flm-3* plants were indistinguishable from *crn,* consistent with the role of the FLM-β in stabilizing SVP (Figures 1C-E). These results support that the *FLM*/*SVP* module may contribute to the restoration of normal flower formation to *crn* at elevated temperatures.

We next asked how loss of either *SVP* or *ELF3*, two unrelated transcriptional regulators, could both restore primordia formation in *crn* plants. While ELF3 plays a role in auxin-mediated thermomorphogenesis, ELF3 is also a repressor of floral transition as part of the “evening complex” that entrains flowering time to circadian inputs^16,27,28^. A complex array of repressors and activators regulates the floral transition, each functioning in distinct and sometimes overlapping circuits^29^. As either *svp* or *elf3* restores flower formation to *crn* at cooler temperatures, we first asked if loss of floral transition repressors in multiple floral transition pathways could similarly restore flower primordia outgrowth to *crn.* We therefore expanded our analysis to the floral transition repressors *PHYTOCHROME B* (*PHYB,* photoperiod pathway repressor), *DELLA* family transcriptional regulators (gibberellic acid (GA) pathway repressor), and the floral transition activator, *CONSTANS (CO,* photoperiod and circadian clock pathway activator)^30–34^. Indeed, loss of either *PHYB* or the *DELLAs*, or overexpression of *CO*, restored flower primordia formation to *crn* mutants at cool and ambient (22°C) temperatures, with *elf3, svp,* and *35S::CO* displaying near complete restoration of primordia outgrowth, and *phyB* and the *della* pentuple mutants (*dellaP*) having an intermediate and temperature-dependent restoration of flower primordia outgrowth (Figures 2A-F, Figures S2A-B). Additionally, we accelerated floral transition by germinating *crn* seedlings on gibberellin (GA_4_)-containing media until 11 days after germination (DAG) and then transferring to soil. This treatment also restored flower formation to *crn* at ambient temperatures (Figure S2C). At cooler temperatures, additional GA_4_ sprays twice weekly until floral transition were necessary to restore flower formation (Figures 2G-H). Collectively, these results indicate that diverse genetic or pharmacological approaches to accelerating floral transition are sufficient to restore primordia formation to *clv* mutants in cooler environments.

**Figure 2:**
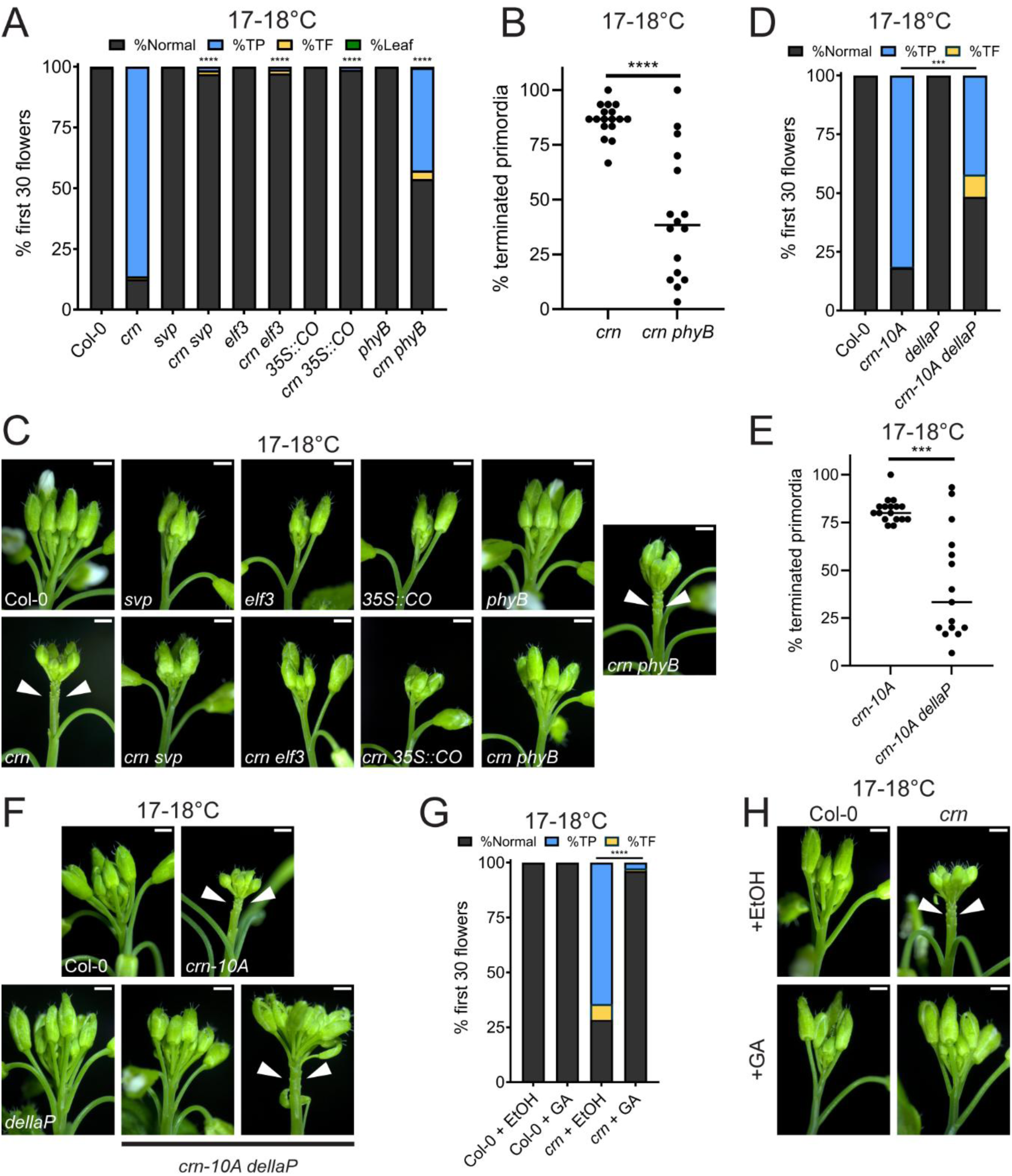
Acceleration of the floral transition restores primordia formation to *crn* mutants. (A) Quantification of floral primordia termination in Col-0 (n=15), *crn* (n=17), *svp* (n=17), *crn svp* (n=13), *elf3* (n=17), *crn elf3* (n=17), *35S::CO* (n=17), *crn 35S::CO* (n=13), *phyB* (n=16), and *crn phyB* (n=16) grown at 17-18°C. Statistical significance of a genotype compared to *crn* is represented on the graph with asterisks. (B) %Terminated primordia for each of the *crn* and *crn phyB* individuals summarized in (A). Line represents the median. (C) Representative inflorescence images of genotypes quantified in (A). (D) Quantification of floral primordia termination in Col-0 (n=18), *crn-10A* (n=17), *dellaP* (n=18), and *crn-10A dellaP* (n=15) grown at 17-18°C. (E) %Terminated primordia for each of the *crn-10A* and *crn-10A dellaP* individuals summarized in (D). Line represents the median. (F) Representative inflorescence images of genotypes quantified in (D). (G) Quantification and (H) representative images of floral primordia termination in Col-0 and *crn* plants germinated on 10µM GA_4_ or ethanol (EtOH) containing plates for 11DAG and sprayed twice weekly with 10µM GA_4_ or ethanol solution until flowering. Sample sizes are Col-0 + EtOH (n=17), Col-0 + GA (n=17), *crn* + EtOH (n=17), and *crn* + GA (n=18.) Statistical significance between %terminated primordia in indicated genotypes were calculated with multiple Mann-Whitney tests using the Holm-Šídák method for correcting multiple comparisons. Significance is represented on the graph using asterisks (p < 0.05 *; p < 0.01 **; p < 0.001 ***; p < 0.0001****.) In (C), (F), and (H), terminated primordia are indicated with arrowheads, and all scale bars are 1mm.

### *FT* buffers the loss of CLAVATA signaling during flower primordia formation

As accelerating floral transition through multiple pathways restores flower outgrowth to *crn* plants, we sought to identify a common effector of these diverse pathways that could mediate this. During the floral transition, many environmentally sensitive transcription factors activate the expression of *FT*, the main component of florigen in *Arabidopsis*, in leaves^35^. FT is then transported through the phloem to the SAM, where it induces floral transition^36–38^. *ELF3*, *SVP*, *DELLAs*, *PHYB*, and *CO* all modulate *FT* transcription in leaves to coordinate the timing of floral transition in response to diverse cues^22,27,30,33,39^. Therefore, we first confirmed that *FT* expression was increased in these genetic backgrounds when combined with *crn* mutations. We detected an increase in *FT* transcript levels in *crn elf3, crn svp, crn phyB,* and *crn 35S::CO* seedlings relative to both 11d Col-0 and *crn* seedlings grown in continuous light at ambient temperatures, consistent with previously published results (Figure S2E)^40–43^. We then asked whether *FT* in these backgrounds is necessary for the restoration of flower production by introducing a null *ft* allele into the appropriate *crn* double mutant backgrounds. Indeed, *crn svp ft, crn elf3 ft, crn phyB ft,* and *crn 35S::CO ft* displayed similar rates of primordia termination as *crn* plants, indicating that increased *FT* expression is necessary for the restoration of primordia formation in *crn* plants (Figures 3A-B). We next asked whether increased *FT* levels are sufficient to restore primordia outgrowth to *crn.* We therefore attempted to complement primordia defects in *crn ft* by expressing *FT* from either the phloem companion cell specific promoter (*pSUC2::FT*) or a SAM-active promoter (*pKNAT1::FT*)^44,45^. In both cases, this resulted in the restoration of normal flower formation in *crn* plants in cooler temperatures, indicating that distal expression of *FT* in phloem is sufficient to elicit SAM primordia, mirroring the mobile signaling functions of FT during floral transition (Figures 3C-D). Collectively, these results indicate that increased *FT* levels are necessary and sufficient to bypass the loss of CLEp signaling during flower formation in cooler environments and can do so as a mobile signal originating from phloem.

**Figure 3:**
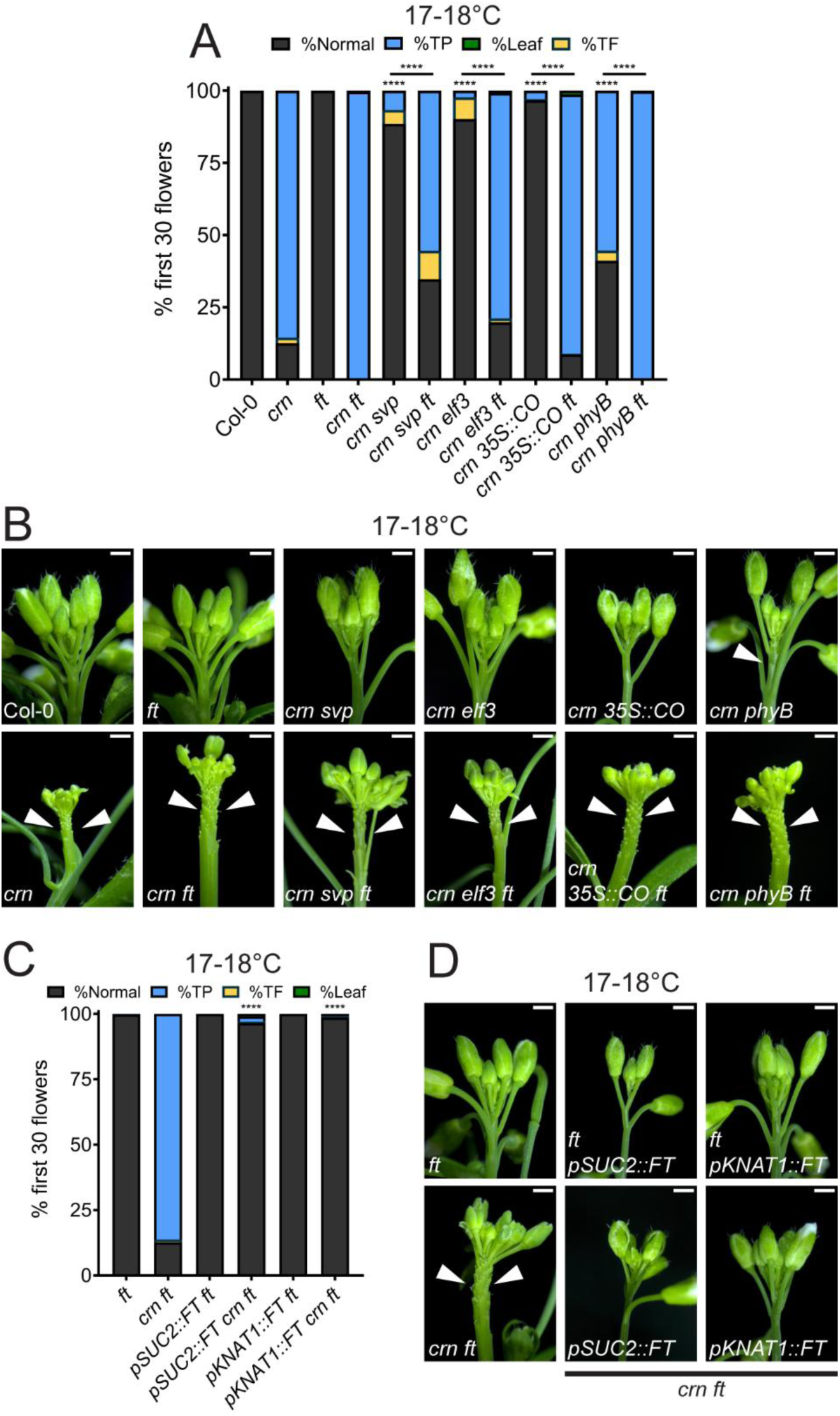
Florigen buffers the loss of CLAVATA signaling during flower primordia formation. (A) Quantification and (B) representative inflorescence images of terminated floral primordia in Col-0 (n=18), *crn* (n=17), *ft* (n=18), *crn ft* (n=18), *crn svp* (n=16), *crn svp ft* (n=17), *crn elf3* (n=16), *crn elf3 ft* (n=18), *crn 35S::CO* (n=17), *crn 35S::CO ft* (n=18), *crn phyB* (n=18), and *crn phyB ft* (n=18) grown at 17-18°C. Statistical comparisons with *crn* are indicated above relevant genotypes. Other pairwise comparisons are indicated on the graph. (C) Quantification and (D) representative images of terminated floral primordia in *ft* (n=17), *crn ft* (n=18), *pSUC2::FT ft* (n=17), *pSUC2::FT crn ft* (n=16), *pKNAT1::FT ft* (n=18), and *pKNAT1::FT crn ft* (n=16) grown at 17-18°C. Statistical comparisons with *crn ft* are indicated above relevant genotypes. Statistical significance between %terminated primordia in indicated genotypes were calculated for (A) and (C) with multiple Mann-Whitney tests using the Holm-Šídák method for correcting multiple comparisons. Significance is represented on the graph using asterisks (p < 0.05 *; p < 0.01 **; p < 0.001 ***; p < 0.0001****.) Arrowheads in (B) and (D) indicate terminated floral primordia. All scale bars are 1mm.

Aside from regulating *FT* expression, SVP also blocks the floral transition by repressing expression of GA biosynthesis genes, and previous work showed that increased GA levels in *svp* mutants accelerate floral transition^46^. As such, we asked whether the restoration of primordia outgrowth to *crn svp* required GA in addition to *FT*. To do this, we genetically abolished all GA biosynthesis in *crn svp* by mutating *GA REQUIRING1* (*GA1*). As *ga1* mutants are unable to germinate without exogenous GA supplementation, we imbibed all genotypes in GA_4_ for 24 hours after the transition to light before plating on normal media, a strategy we adapted from Silverstone *et al*^47^. This treatment did not restore primordia formation to *crn* plants but permitted germination of *ga1* mutants. We observed normal flower production in *crn svp ga1* plants grown at cooler temperatures, comparable to *crn svp* double mutants (Figures S3A-B). As such, increased GA biosynthesis in *svp* mutants is dispensable for the restoration of flower primordia production in *crn svp* plants, contrasting with the clear requirement for *FT* (Figures 3A-B). As GA promotes the floral transition through *FT*-dependent and *FT*-independent mechanisms, we also asked whether restoration of primordia outgrowth to *crn* by GA_4_ treatment was *FT*-dependent^30,48,49^. GA_4_ treatment significantly restores flower formation to *crn ft* plants at cooler temperatures, indicating that *FT* is dispensable for GA-mediated restoration of primordia outgrowth in *crn* plants (Figures S3C-E). Consistent with this, *FT* levels were not upregulated in 11d *crn-10A dellaP* seedlings grown at ambient temperature in continuous light conditions (Figure S2E). In summary, these results show that the acceleration of floral phase transition through diverse mechanisms is sufficient to restore primordia formation in *crn* plants, with *FT* requirements mirroring those in floral transition pathways^30,48,49^.

### *FT* promotes floral primordia formation through SAM expressed transcriptional regulators

During the floral transition, FT interacts with the basic leucine zipper (bZIP) transcription factor FD to activate the floral fate transcriptional program^50–52^. To test if the restoration of primordia formation in *crn* plants by increased *FT* requires *FD*, we generated *crn svp fd* mutants. These plants displayed rates of terminated primordia similar to *crn,* consistent with a role for FD-dependent transcriptional activity in promoting flower primordia restoration in *crn* mutants (Figures S4A-B). FT competes with its paralog TERMINAL FLOWER1 (TFL1), which blocks floral transition, for FD binding^52–54^. *TFL1* expression levels are slightly elevated in *crn* SAMs in our RNA-Seq dataset, raising the possibility that ectopic *TFL1* might block formation of FD-FT complexes to prevent normal flower primordia formation in *crn* plants (Figure S4C). To test this, we generated *crn tfl1* double mutants. While we could not interpret the effect of the strong *tfl1-1* allele on *crn* primordia termination due to the nature of the terminal flower phenotype associated with *tfl1-1* mutants, using the weak *tfl1-11* allele we observed a minor decrease in primordia termination in *crn tfl1-11* compared to *crn* single mutants (Figures S4D-E). These results suggest that ectopic *TFL1* does not substantially contribute to primordia termination in *crn* SAMs. Our results indicate that elevated *FT* levels bypass the requirement for CLEp signaling during flower formation in cool environments in an FD-dependent manner.

As part of the later floral transition phase, FT-FD promote expression of multiple floral meristem identity factors that reinforce floral fate in developing primordia (Figure 4A)^42,50–52,55,56^. As such, we wondered if floral meristem identity targets were genetically required for flower restoration downstream of *FT*. To test this, we mutated four floral identity factors—*LEAFY* (*LFY*)*, APELATA1* (*AP1*)*, SUPPRESSOR OF OVEREXPRESSION OF CONSTANS1* (*SOC1*), and *AGAMOUS-LIKE24* (*AGL24*)—individually in the *crn svp* background, except for *clv2 svp lfy* due to genetic linkage between *CRN* and *LFY*. As *lfy* and *ap1* plants produce flowers with floral identity or floral reversion defects, we classified all lateral organ attempts as either a terminated primordia or “outgrown organ.” At cooler temperatures, *crn svp ap1* and *crn svp agl24* plants displayed slightly increased rates of primordia termination compared to *crn svp*, while *crn svp soc1* plants were similar to *crn svp* mutants. In contrast, *clv2 svp lfy* mutants displayed levels of organ termination similar to *clv2* at cooler temperatures (Figures 4B-C). At ambient temperatures, *crn svp ap1* and *crn svp agl24* plants showed normal flower formation, *crn svp soc1* plants displayed a modest increase in primordia termination, and *clv2 svp lfy* plants again showed similar rates of primordia termination as *clv2* (Figure S2D). Together, these results suggest that *LFY* plays a dominant role downstream of *FT* to promote flower primordia restoration to *crn* plants in cool environments, though the other floral integrators likely contribute quantitatively to *FT-*mediated primordia outgrowth redundantly with *LFY*.

**Figure 4:**
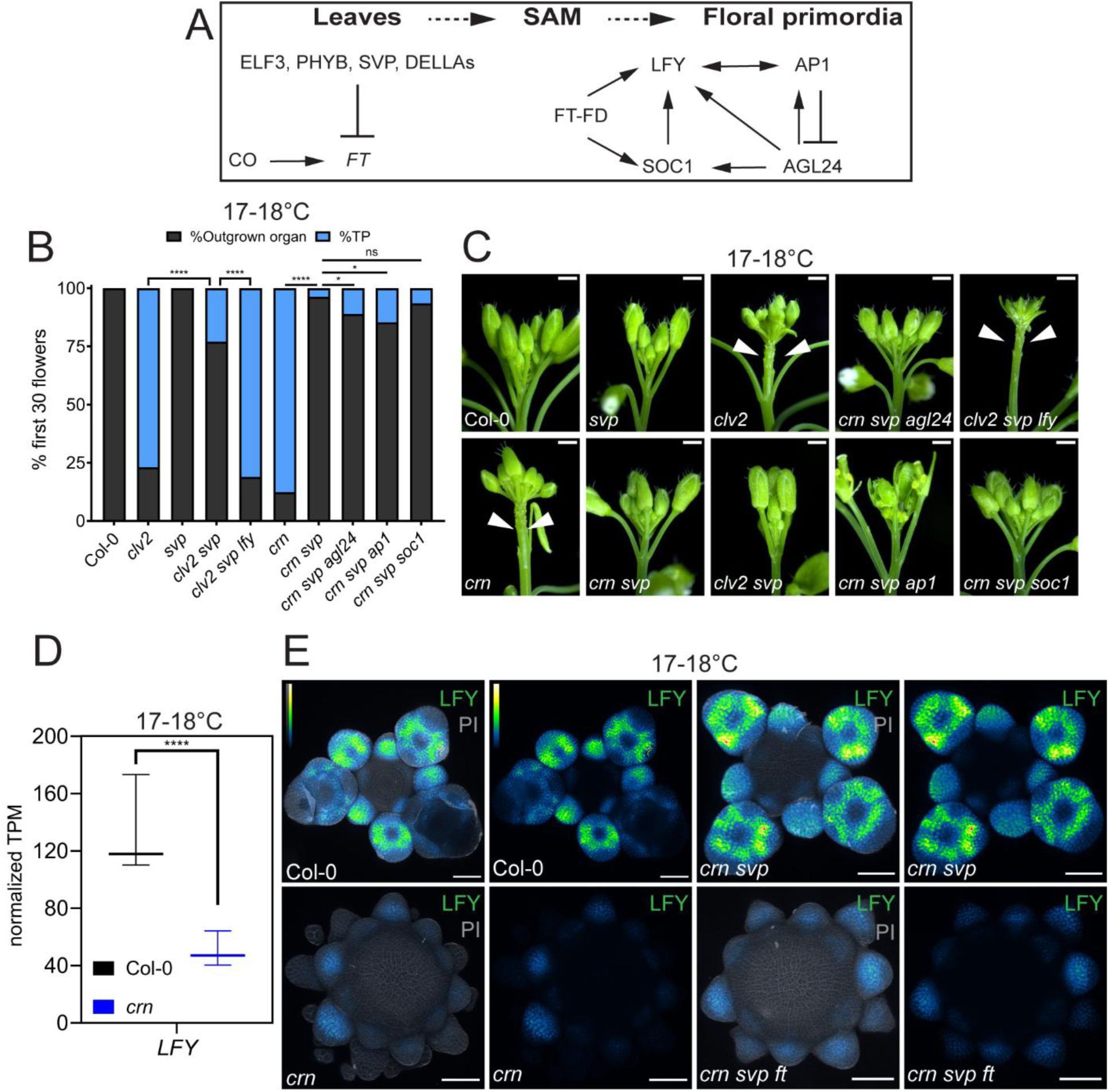
Florigen promotes floral primordia formation through SAM expressed transcriptional regulators. (A) Schematic of gene interactions during the floral transition and activation of floral identity in incipient floral primordia. (B) Quantification and (C) representative inflorescence images of floral primordia termination in Col-0 (n=18), *clv2* (n=15), *svp* (n=16), *clv2 svp* (n=8), *clv2 svp lfy* (n=12), *crn* (n=18), *crn svp* (n=16), *crn svp agl24* (n=16), *crn svp ap1* (n=18), and *crn svp soc1* (n=17) grown at 17-18°C. All outgrown organs (normal flowers, flowers with identity defects, leaves, terminated flowers) were pooled into an “outgrown organ” category distinct from terminated primordia. Statistical comparisons are indicated on the graph. (D) Expression levels of *LFY* from RNA-Seq analysis in Col-0 and *crn* SAMs grown at 17-18°C. Normalized transcripts per million (TPM) summed for *LFY* for three biological replicates are plotted from Kallisto/Sleuth RNA-Seq analysis. The line indicates the median. P values for the three isoforms of *SVP* were p=0.000171 and p=0.000934. (E) Maximum intensity projection of *pLFY::LGY-GFP* (GFP signal is color-coded for intensity using the BlueToYellow LUT) with PI (Gray) and *pLFY::LFY-GFP* alone in Col-0 (n=18), *crn* (n=11), *crn svp* (n=18), and *crn svp ft* (n=8) SAMs grown at 17-18°C. Statistical significance between %terminated primordia in indicated genotypes were calculated for (B) with multiple Mann-Whitney tests using the Holm-Šídák method for correcting multiple comparisons. Significance is represented on the graph using asterisks (p < 0.05 *; p < 0.01 **; p < 0.001 ***; p < 0.0001****.) Arrowheads in (C) indicate terminated floral primordia. Scale bars are 1 mm in (C) and 50 µM (E).

To further test this hypothesis, we examined *pLFY::LFY-GFP* accumulation in incipient primordia. We observed a decrease in *pLFY::LFY-GFP* accumulation in aborted primordia in *crn,* consistent with a loss of flower formation and reduced *LFY* expression levels in our RNA-Seq dataset (Figures 4D-E). Accelerating the floral transition in *crn svp* restored *pLFY::LFY-GFP* accumulation in primordia to wild type levels. This increase was dependent on *FT*, as *pLFY::LFY-GFP* levels were reduced to *crn* levels in *crn svp ft* (Figure4E). Collectively, these results support a role for *LFY* downstream of *FT* during the canalization of flower production, consistent with recent evidence demonstrating that FT-FD binds directly to the *LFY* promoter to activate its expression^52,57^. As such, in cooler environments, loss of CLEp signaling in floral primordia formation can be buffered by increased *FT* expression which acts via a FD/FT-LFY transcriptional circuit in SAMs.

### Thermomorphogenesis-mediated canalization of flower production requires *FT*/*TSF*

Our original observation that elevated temperatures can restore floral primordia formation and outgrowth to *clv* mutants reveals that thermomorphogenic responses canalize floral development. The mechanistic basis of this is unknown. Our work reveals that increased *FT* levels are sufficient to restore primordia formation to *crn* plants in cooler environments. As high temperatures also accelerate floral transition by inducing *FT* expression, we asked whether thermomorphogenesis-induced canalization of primordia formation in SAMs is an *FT* and *LFY-*dependent process^58^. Indeed, *crn ft* plants displayed terminated primordia in the heat, indicating that *FT* mediates heat-induced flower primordia restoration in *crn* (Figures 5A-B). The *FT* paralog *TSF* can partially compensate for the loss of *FT* during floral transition^59,60^. Consistent with this, we observed increased primordia termination in heat-grown *crn tsf ft* plants relative to *crn ft* plants (Figures 5A-B). FT and TSF both interact with FD to induce the floral fate transcriptional program^42^. Heat- grown *crn fd* plants displayed lower levels of terminated primordia relative to *crn tsf ft* plants (Figures 5A-B), possibly due to the partial redundancy of *FD* with the related bZIP transcription factor *FD PARALOG (FDP)*^61^. At a minimum, our analysis reveals that primordia formation during thermomorphogenesis is canalized in a *FT*/*TSF*-dependent manner, and most likely acts through FD and associated factors.

**Figure 5:**
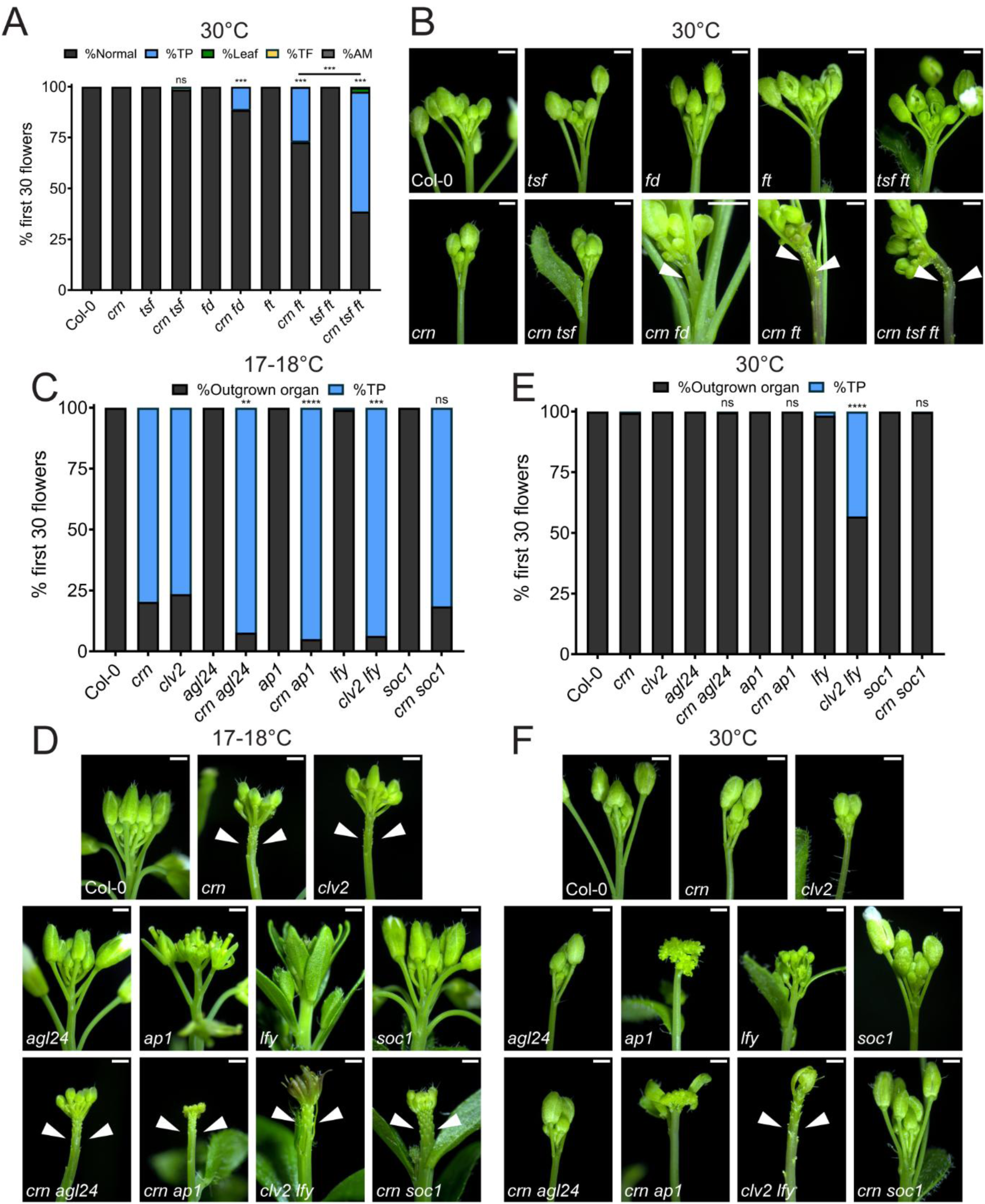
Heat-mediated canalization of flower production requires florigen. (A) Quantification and (B) representative inflorescence images of terminated floral primordia in Col-0 (n=19), *crn* (n=17), *tsf* (n=13), *crn tsf* (n=16), *ft* (n=18), *crn ft* (n=18), *fd* (n=18), *crn fd* (n=18), *tsf ft* (n=4), and *crn tsf ft* (n=20) plants grown at 30°C. Axillary meristems (AM) are shown in gray. Statistical comparisons with *crn* are indicated above the relevant genotypes. Other comparisons are indicated on the graph. Note increased magnification in *crn fd* to illustrate terminated primordia. (C) Quantification and (D) representative inflorescence images of terminated floral primordia in Col-0 (n=18), *crn* (n=18), *clv2* (n=18), *agl24* (n=18), *crn agl24* (n=16), *ap1* (n=18), *crn ap1* (n=18), *lfy* (n=18), *clv2 lfy* (n=14), *soc1* (n=18), *crn soc1* (n=18) plants grown at 17-18°C. Statistical comparisons with *crn* or *clv2* are noted over the relevant genotypes. (E) Quantification and (F) representative inflorescence images of terminated floral primordia in Col-0 (n=18), *crn* (n=16), *clv2* (n=17), *agl24* (n=15), *crn agl24* (n=15), *ap1* (n=13), *crn ap1* (n=18), *lfy* (n=17), *clv2 lfy* (n=12), *soc1* (n=17), *crn soc1* (n=18) plants grown at 17-18°C. Statistical comparisons with *crn* or *clv2* are noted over the relevant genotypes. Statistical significance between %terminated primordia in indicated genotypes were calculated for (A, C, E) with multiple Mann-Whitney tests using the Holm-Šídák method for correcting multiple comparisons. Significance is represented on the graph using asterisks (p < 0.05 *; p < 0.01 **; p < 0.001 ***; p < 0.0001****.) Arrowheads in (B), (D), and (F) indicate terminated floral primordia. All scale bars are 1mm.

To assess whether heat-mediated primordia restoration requires *LFY*, or other floral identity factors acting downstream of *FT*, we generated double mutants of either *clv2* or *crn* and mutations in different individual floral identity regulators, again utilizing both *clv* mutants due to genetic linkage between *CRN* and *LFY*. As *clv2 lfy* and *crn ap1* plants display flowers with various identity defects or characteristics with inflorescence reversion, we again categorized all lateral organs attempts as either an “outgrown organ” or “terminated primordia.”^62,63^ At cool temperatures, *agl24, ap1,* and *lfy* modestly enhanced terminated primordia numbers in *crn*/*clv2,* while *soc1* had no effect (Figures 5C-D). In the heat, *clv2 lfy* plants displayed numerous terminated primordia, while *crn soc1, crn agl24,* and *crn ap1* had normal organ formation (Figures 5E-F). As such, thermomorphogenesis acts to canalize floral primordia formation through *FT*/*TSF* and floral identity factors, with *LFY* playing a dominant role.

As *FT*/*TSF* are necessary for heat-mediated floral primordia formation, we next asked which repressors of *FT* might act as the thermosensors in this process. ELF3 and SVP indirectly and directly repress *FT,* respectively, and are inactivated by heat, through protein condensation and proteasomal degradation respectively^18,23,26,27^. The FLM*-*β isoform of FLM, which accumulates at cooler temperatures, stabilizes SVP to repress *FT* transcription^23,26^. Despite this, *FLM-β-GFP crn flm-3* plants make normal flowers at high temperatures, indicating that blocking SVP degradation alone is not sufficient to prevent heat-mediated primordia outgrowth (Figures S5A-C). As such, we expressed an ELF3 variant (*ELF3-BdPrD*), which is insensitive to heat-mediated inactivation, under the native *ELF3* promoter and introduced it into *FLM-β-GFP crn flm-3* plants^18^. Despite this, *crn flm-3 FLM-β-GFP elf3 ELF3-BdPrD-GFP* plants displayed normal flower formation in heat, indicating that simultaneously blocking ELF3 and SVP thermosensing is insufficient to prevent heat-induced flower formation (Figures S5A-C). This suggests the existence of additional *FT*- regulating thermosensors acting in heat-mediated primordia formation.

### CLAVATA and florigen signaling synergize to canalize flower formation across broad temperature environments

Overall, our results show that the thermomorphogenically accelerated floral transition induction also canalizes flower primordia production, revealing a novel role for *FT* in promoting floral primordia initiation and outgrowth. We therefore asked whether CLEp signaling and *FT/TSF* promote primordia outgrowth in overlapping or independent pathways. We first took advantage of the negative regulator of CLEp signaling *POLTERGEIST* (*POL*), a phosphatase which dephosphorylates the CLV1 receptor to dampen signaling^64^. Previously, we showed that mutating *POL* restores floral primordia outgrowth to *crn* due to ectopic activation of CLV1^12,13^. To ask whether CLV1*-*mediated restoration of flower formation requires *FT*, we generated *crn pol ft* mutants. At cool temperatures, these plants displayed normal primordia outgrowth, indicating that CLEp signaling promotes flower outgrowth independently of *FT* (Figures 6A-B). Further supporting this hypothesis is our observation that *FT* expression levels in 11d *crn pol* seedlings grown in continuous light conditions at ambient temperature are comparable to Col-0 despite the rescue of flower formation (Figure S2E). Altogether, these data suggest that florigen and CLEp signaling operate in parallel to promote flower formation.

**Figure 6.**
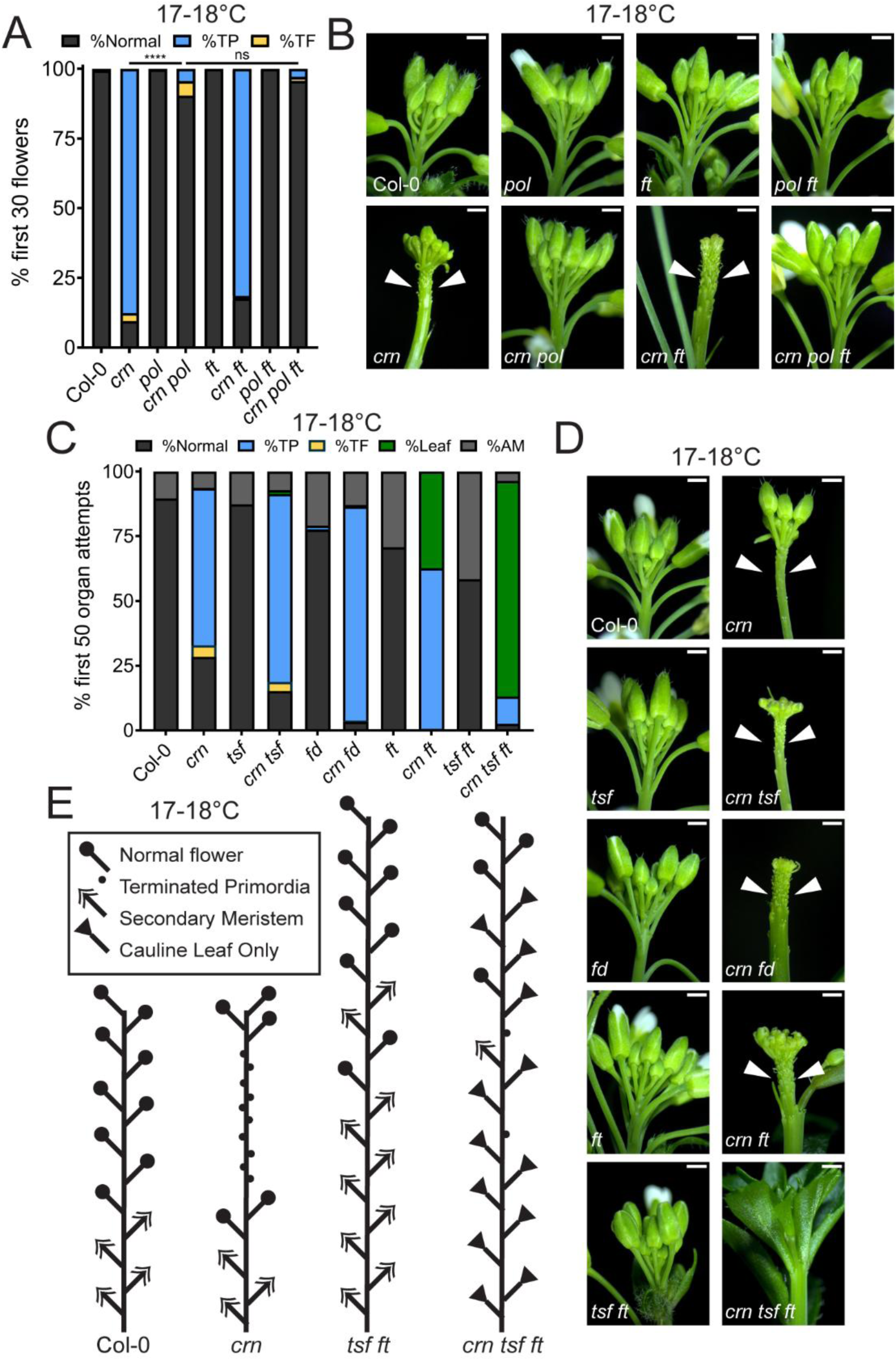
CLAVATA and florigen signaling synergize to canalize flower formation across broad temperature environments. (A) Quantification and (B) representative inflorescence images of floral primordia termination in Col-0 (n=18), *crn* (n=18), *pol* (n=16), *crn pol* (n=18), *ft* (n=16), *crn ft* (n=18), *pol ft* (n=15), and *crn pol ft* (n=16) plants grown at 17-18°C. Statistical comparisons indicated on the graph. (C) Quantification and (D) representative inflorescence images of the first fifty organ attempts in Col-0 (n=16), *crn* (n=16), *tsf* (n=13), *crn tsf* (n=16), *fd* (n=18), *crn fd* (n=17), *ft* (n=15), *crn ft* (n=14), *tsf ft* (n=18), and *crn tsf ft* (n=17) plants grown at 17-18°C. Organ attempts were classified as normal (black), terminated primordia (blue), terminated flowers (yellow), leaves (green), or axillary meristems (AM, gray.) (E) Schematic of Col-0, *crn, tsf ft,* and *crn tsf ft* organ attempts at end-of-life when grown at 17-18°C. Statistical significance between %terminated primordia in indicated genotypes were calculated for (A) with multiple Mann-Whitney tests using the Holm-Šídák method for correcting multiple comparisons. Significance is represented on the graph using asterisks (p < 0.05 *; p < 0.01 **; p < 0.001 ***; p < 0.0001****.) All scale bars are 1mm. Arrowheads indicate terminated primordia.

Even though our data reveals that *FT*/*TSF* promotes lateral organ outgrowth, we observed normal flower formation in *tsf ft* plants in the cooler temperatures, consistent with previously published results (Figures 6C-D)^42^. This suggests the presence of an additional pathway that bypasses the requirement for *FT*/*TSF* to promote primordia formation in cooler temperatures. To ask whether CLEp signaling facilitates this canalization, we inspected *crn tsf ft* triple mutants growing at cooler temperatures. We characterized lateral organ attempts as normal flowers, terminated primordia, terminated flowers, cauline leaves, or axillary meristems (AMs, branch subtended by cauline leaf). In *crn tsf ft* plants, most lateral organs in the first fifty attempts were cauline leaves, with some terminated primordia and terminated flowers. Rarely, flowers (with or without inflorescence reversion) or normal branches were produced (Figures 6C-D). After the first fifty organ attempts, nearing when Col-0 plants senesce, *crn tsf ft* plant continued growth and then stochastically produce flowers intermixed with cauline leaves, terminated primordia, and terminated flowers (Figure 6E, FigureS6). As such, these results indicate that CLEp signaling and *FT*/*TSF* synergize to canalize flower production across a broad range of environmentally relevant temperatures, with CLEp signaling being essential in cooler environments, and both pathways contributing to primordia formation in heat (Figures 5A-B).

## Discussion

How organisms maintain essential developmental pathways in the face of fluctuating environmental conditions is an outstanding question in biology. For plants, ensuring robust reproduction and therefore reliable agricultural yields across broad temperature ranges is an outstanding goal, but few mechanisms that promote robust flower formation during environmental perturbation are described. Here, our work reveals that flower formation is canalized by CLEp signaling and environmentally mediated titration of florigen levels across broad temperatures ranges (Figure 7). At cooler temperatures, CLEp signaling plays a more dominant role in promoting flower production, as floral primordia terminate in cool grown *crn* mutants. However, our data shows that elevated *FT*/*TSF* expression levels bypass the requirement for CLEp signaling at any temperature; *crn tsf ft* triple mutants show a total loss of flower formation within the normal *Arabidopsis* lifespan at cool temperatures and have strongly compromised flower production at elevated temperatures. Curiously, while *TSF* plays a minor role promoting floral transition, our data suggest that it contributes to the canalization of flower production^42,59^. While *crn ft* plants can produce flowers after the termination phase, loss of *tsf* in this background abolishes flower production until the end of life, indicating that *TSF* is likely responsible for the flower formation observed in *crn ft.* This data highlights the role of paralogs in facilitating canalization of important traits^65^. The regulation of floral transition is highly complex. Based on our results here, we predict that other environmental conditions or genetic perturbations that also elevate *FT*/*TSF* expression levels would also buffer against loss of CLEp signaling in flower primordia formation. This could explain why flower formation in *crn* is so strongly canalized in elevated temperatures, and our data suggests that additional thermosensors besides SVP and ELF3, can upregulate *FT*/*TSF* expression during thermomorphogenesis to allow flower formation. Based on our analysis here of *crn phyb* and *crn phyb ft* mutants, and its known temperature sensitivity, phyB is a likely candidate^66–68^, although others may exist.

**Figure 7.**
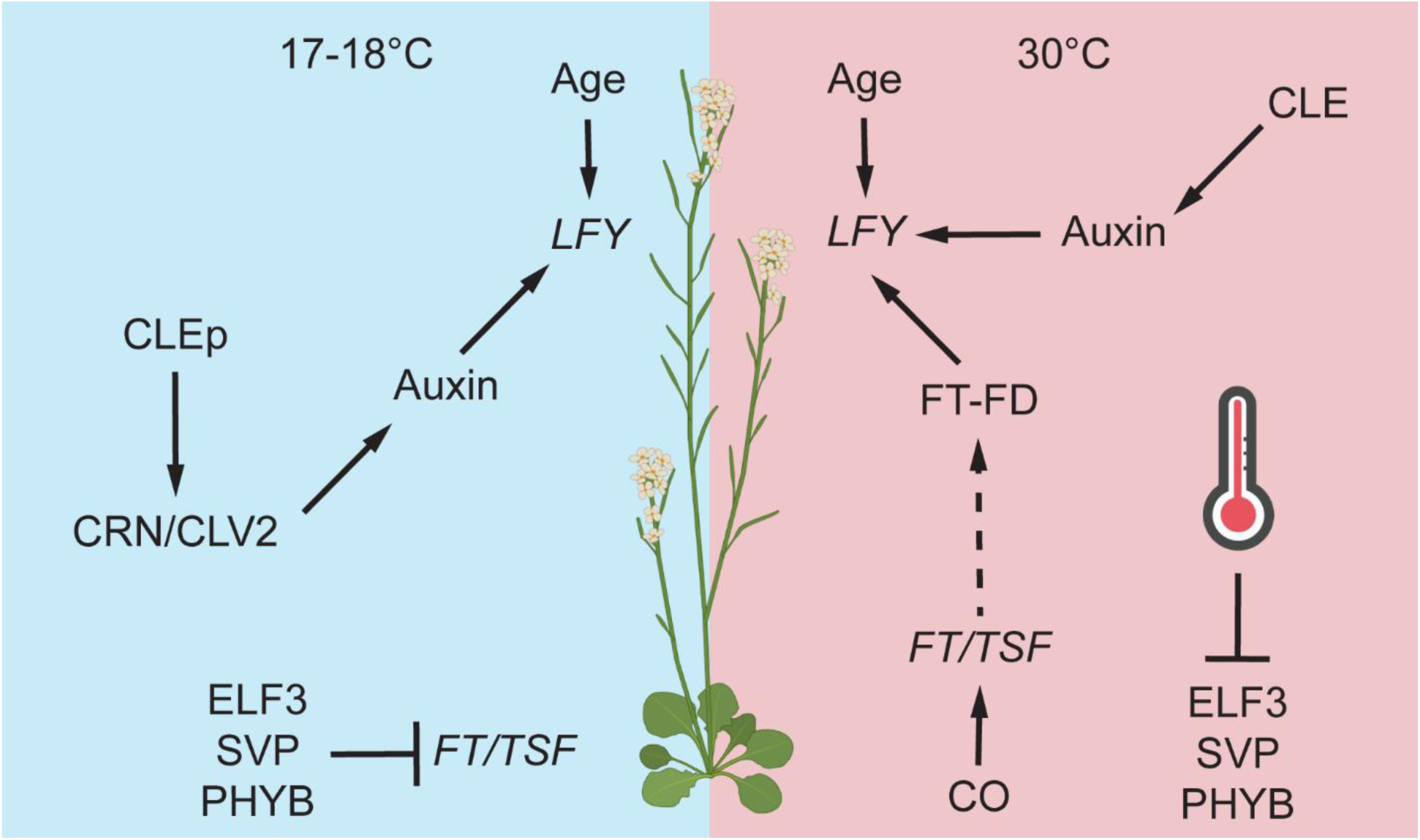
Canalization of flower production across thermal environments requires Florigen and CLAVATA signaling. Schematic of gene interactions promoting floral primordia formation and outgrowth across temperature regimes.

In cooler environments, *crn* plants produce 2-5 normal flowers before entering the terminated primordia phase. As no early flowers are formed in *crn tsf ft* plants, this implies that the first few normal flowers formed in *crn* plants before the termination phase are due to the *FT*/*TSF*-induced floral transition. Following this switch activity, *crn* plants abort later floral primordia formation. As such, there is a temporal control of normal flower production in plants grown at cooler temperatures, with *FT*/*TSF* functioning early, and CLEp signaling being essential later. Increasing *FT*/*TSF* levels, either through heat or inactivation of *FT* repressors, eliminates the terminated primordia phase in *crn* plants. This work reveals that *FT*/*TSF* can act as both a phase switch and a sustained signal molecule promoting floral primordia initiation and outgrowth in SAM morphogenesis. This sustained activity role mirrors the function of the *FT-FD* module in preventing floral reversion^69,70^. Our work shows that the function of florigen following the floral transition extends to the initiation and outgrowth of early floral primordia before the floral meristem itself is established.

Here, we show that this thermomorphogenic increase in *FT*/*TSF* expression promotes flower primordia formation, but that flower primordia formation is not sensitized to ELF3-mediated auxin biosynthesis circuits, even though *ELF3* is expressed in SAMs in three published SAM RNA-Seq data sets, including SAM cell specific transcriptomes ^13,71,72^. Previously we showed that auxin is essential for heat-mediated primordia formation, as mutating two *YUC*s that are integral to floral primordia formation—*YUC1* and *YUC4—*in the *clv2* background resulted in pin inflorescences in heat^13,73^. However, unlike *YUC8*, *YUC1* and *YUC4* do not contain *PIF* binding sites in their promoters and are not heat-sensitive PIF4 targets during ELF3-dependent thermomorphogenesis in other tissues^19^. This suggests that thermomorphogenic transcriptional and subsequent growth responses are specialized to tissue type, expanding on previous observations^6^. Our data also shows that thermomorphogenesis controls SAM patterning through transcriptional changes and mobile signal production in leaves. Forced auxin signaling in stem cells can have deleterious impacts on SAM functions^11,74,75^. While it is unclear if heat increases auxin biosynthesis rates in SAMs at high temperatures, we clearly show that the heat induced *PIF4*-*YUC8* circuit is dispensable for primordia formation in heat (Figure S1). As finely tuned hormone signaling fields in the SAM position new organ primordia in the correct phyllotaxy, tying thermomorphogenic responses in SAMs to distal florigen, and not local *YUC* activation, may be a mechanism to prevent SAM patterning disruptions by temperature-induced hormone changes^76–80^.

After producing aborted floral primordia, *crn* plants at cool temperatures produce normal flowers again. Even *crn tsf ft* plants eventually produce flowers in older plants after extensive organ attempts. Previous work showed that the age and GA floral transition pathways can redundantly activate expression of *LFY* and other floral integrators independent of *FT*/*TSF*, explaining flower formation in *tsf ft*^48,49,81^. Interestingly, a florigen sextuple mutant in which age-dependent floral transition is knocked down through overexpression of *miR156* still produces flowers. The authors of that study speculated the existence of a third pathway promoting floral transition, and our data suggests that CLEp signaling could function as that third pathway during both floral transition and primordia initiation and outgrowth^54^. Furthermore, age-dependent activation of *LFY* expression may explain flower formation during the recovery phase of *crn* plants grown at cool and ambient temperatures^48,81^. Further investigation into the interactions between these pathways, as well as elucidating the mechanisms by which CLEp signaling itself promotes primordia formation, will clarify our understanding of the molecular mechanisms underlying canalized flower production. Collectively, our work shows that environmental titration of *FT*/*TSF* levels buffers flower production against stochastic mutations in CLEp signaling machinery to ensure primordia formation. Many crop species were bred to have mutations in various CLEp signaling components, as defects in CLEp signaling increase cell proliferation in the floral meristem size and thereby fruit size^82–87^. Additionally, floral transition pathways are a major trait target in many crop plants^88,89^. It is possible that further genetic engineering of CLEp signaling and florigen levels could be used to help buffer crop flower formation against current and future environmental fluctuations.

## Methods and reagents

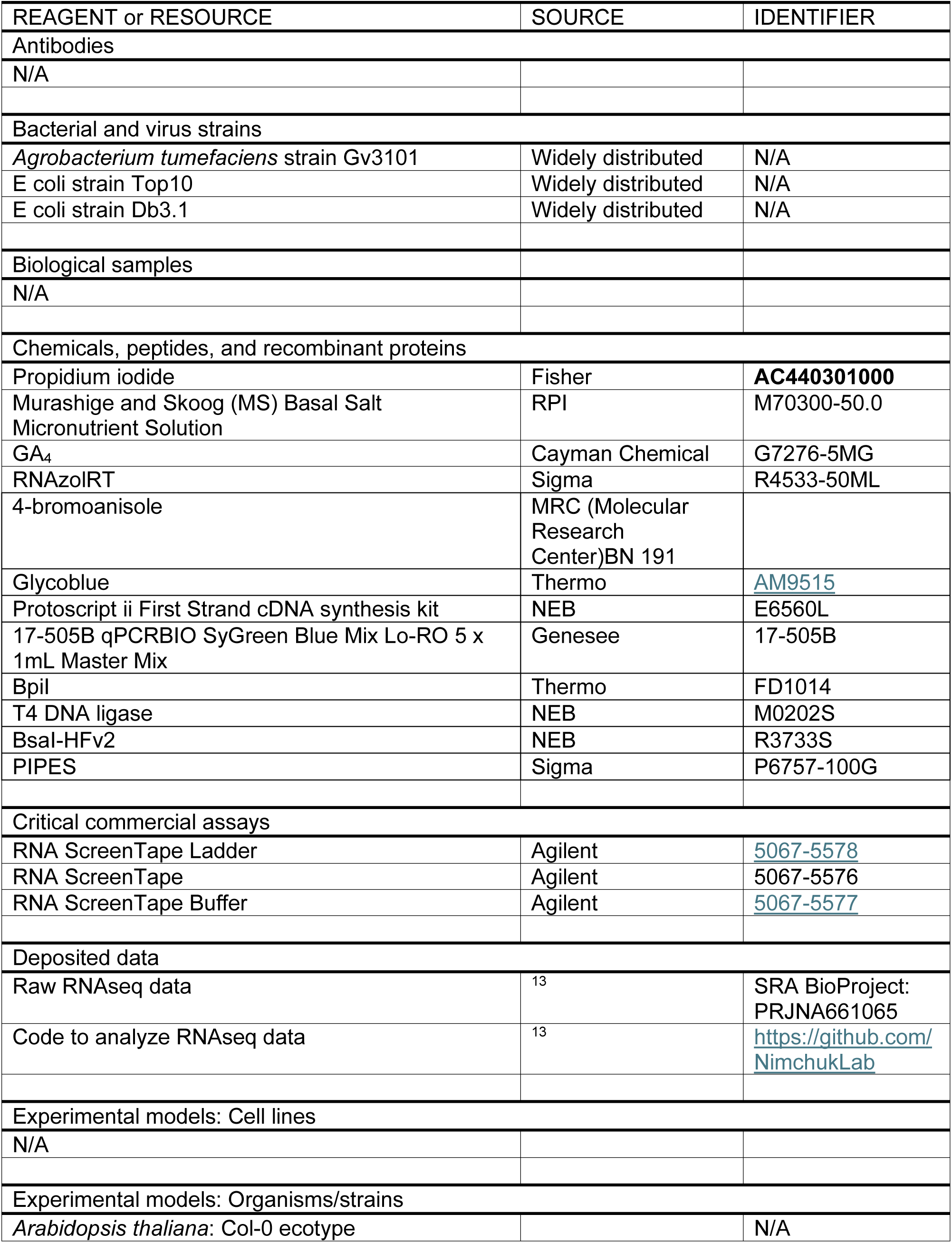
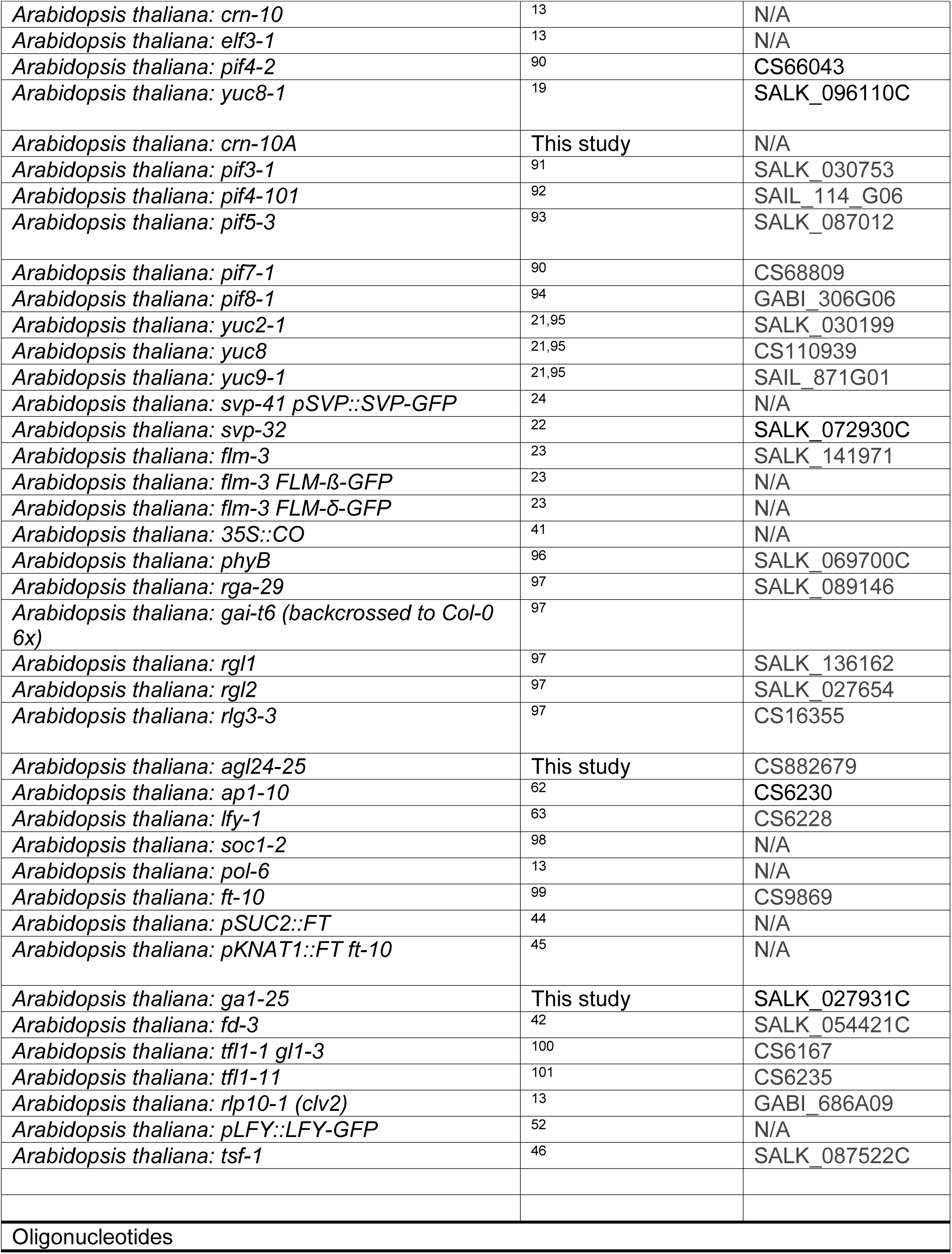
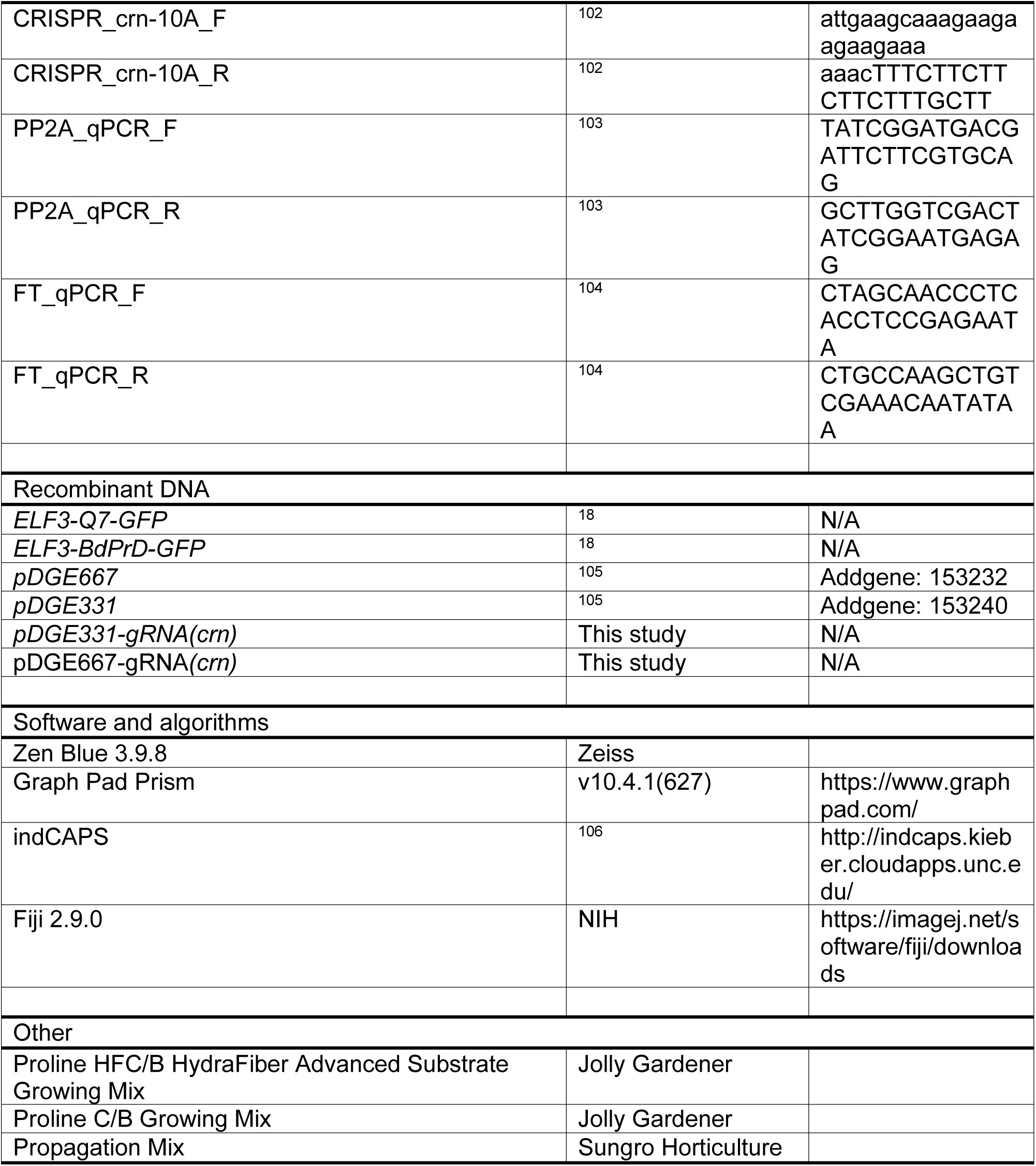

### Lead Contact

Requests for reagents, resources, or information should be directed to and will be fulfilled by the Lead Contact, Zachary Nimchuk.

### Materials Availability

*Arabidopsis* lines generated through this study are freely available to academic researchers through the Lead Contact.

### Experimental Model and Subject Details

The *Arabidopsis thaliana* ecotype Col-0 was used as the primary model system throughout this work.

### Plant growth conditions

After sterilization, seeds were plated on half-strength MS (Murashige-Skoog) media buffered with MES, pH= 5.7. Seeds were stratified at 4°C in the dark for two days and transitioned to a continuous light chamber kept at 22°C. One week after transitioning to light, plants were transplanted to soil (Proline HFC/B, Proline C/B, or propagation mix combined with Perlite and supplemented with recommended levels of Peter’s 20:20:20 [N:P:K]). For T1 experiments and GA_4_ treatment, both controls and T1 seedlings were transplanted at 11DAG instead of 7DAG.

After transplanting, plants were transferred to continuous light growth chambers or a custom-built growth room at the relevant temperature. Assays at 17-18°C took place in one of two Percival growth chambers (9AR-75L3 and AR66L3). Assays at 22°C took place in a custom-built growth room. Assays at 30°C took place in a Percival growth chamber (AR66L3). Experiments comparing the same genotypes at 17-18°C and 30°C were grown in both AR66L3 chambers.

### Plant materials

All mutant alleles in this study are in the Col-0 ecotype. All genotyping primers are listed in TableS1. The following genotypes and reporters have been previously described: *crn-10*^13^*, elf3-1*^13^*, pif4-2*^107^*, yuc8-1*^108^*, pif3-1*^91^*, pif4-101*^92^*, pif5-3*^93^*, pif7-1*^107^*, pif8-1*^94^*, yuc2-1*^21,95^*, yuc8*^21,95^*, yuc9-1*^21,95^*, svp-32*^22^*, flm-3*^23^*, flm-3 FLM-ß-GFP*^23^*, flm-3 FLM-δ-GFP*^23^*, pLFY::LFY-GFP*^52^*, svp-41 pSVP::SVP-GFP*^24^*, 35S::CO*^41^*, phyB*^96^*, rga-29*^97^*, gai-t6* (originally in Ler but backcrossed to Col-0 six times)^97^, *rgl1*^97^*, rgl2*^97^*, rgl3-3*^97^*, ap1-10*^62^*, lfy-1*^63^*, soc1-2*^98^*, pol-6*^13^*, ft-10*^99^*, pSUC2::FT*^44^*, pKNAT1::FT ft*^45^*, fd-3*^42^*, tfl1-1 gl1-3*^100^*, tfl1-11*^101^*, rlp10-1 (clv2)*^13^, and *tsf-1*^46^. An *agl24-25* allele with an insertion in the fourth exon was selected (SALKseq_2434). For *ga1-25,* we used SALK_027931C. This line contains an insertion in the eleventh exon and displays phenotypic traits comparable to previously published *ga1* mutants (necessity for exogenous GA for germination, darker rosette leaves, smaller rosette, delayed flowering, reduced internode elongation, etc.)

*crn-10A pif34578, crn-10A yuc289,* and *crn-10A dellaP* were generated using CRISPR Cas9 (described below) to knock out *crn* in previously generated *pif34578 (pif3-1, pif4-101, pif5-3, pif7-1, pif8-1), yuc289*^21,95^ *(yuc2-1, yuc8, yuc9-1),* and *dellaP*^97^ (*rga-29, gai-t6, rgl1, rgl2, rgl3-3)* mutants. A single *crn-10A* mutant was also generated to use as a control. Several SSLPs on chromosome 1 near the GAI locus were genotyped in *crn-10A dellaP* T2 individuals; all loci genotyped as Col-0 in all T2 individuals tested. Progeny from one of these individuals was bulked to use in phenotyping assays.

### CRISPR mutagenesis of *crn*

To expediently generate higher order knockouts, *crn-10A pif34578, crn10-A yuc289, crn-10A dellaP,* and a *crn-10A* control line were generated using the zCas9i cloning kit. This kit uses an intron-optimized and GFP tagged version of Cas9^105^. The previously published guide sequence aagcaaagaagaagaagaaa for *crn* was used^102^. The guide sequence was cloned into the shuttle vector *pDGE331* using modified Golden Gate with BpiI. Then, the cassette containing the *Arabidopsis* U6 promoter, the guide sequence, and the sgRNA was cloned into *pDGE667* using modified Golden Gate cloning with BsaI-HFv2. This plasmid was transformed into *Agrobacterium* and introduced to *Arabidopsis* using the floral dip method^109^. T1 plants were selected on hygromycin containing media and genotyped for Cas9. Plants positive for Cas9 were sequenced at the 5’ end of the *CRN* locus to characterize editing. Plants with homozygous insertion of a single adenine at the 21^st^ base pair were taken to the next generation. This insertion site is the same as the published *crn-10* allele, though the nucleotide inserted in *crn-10A* is adenine instead of the thymine in *crn-10*. This created an early frame-shift truncated protein of the same length with one amino acid change relative to *crn-10.* T2s were screened for seed coat FASTRED; seeds with no signal were plated and subsequently genotyped as previously described to ensure no Cas9 remained in the genome^105^. The *CRN* locus was sequenced to ensure homozygosity of the correct allele. Seeds from a single T2 plant were bulked and used for experiments.

### GA_4_ treatment

For experiments germinating seedlings on GA_4_ plates, a 10 mM stock solution of Giberellin (GA_4_) (Cayman Chemical) was prepared in 100% Ethanol. This was diluted to a final concentration of 10 µM in ½ MS plates. Vehicle control plates were prepared using an equal volume of ethanol in ½ MS. A 10 µM GA_4_ solution was prepared for spraying by diluting the 10 mM stock in water and 0.02% Silwet. A control solution with equal volume of ethanol was also prepared. This method was adapted from Andrés *et al*.^46^

A protocol for GA_4_ treatment during the first 24 hours after germination was adapted from Silverstone *et al.*^47^ Seeds were sterilized normally and resuspended in a 0.1 mM GA_4_ solution. These were stratified for two days in the dark at 4°C. Then, seeds were transferred to light in a 22°C growth chamber. Following 24 hours of light exposure, seeds were washed four times with water, plated on ½ MS plates, and returned to the growth chamber.

### Generation of transgenic lines

New transgenic lines for T1 experiments were generated by transforming the relevant plasmid to *Agrobacterium* and introducing the construct to *Arabidopsis* using the floral dip method^109^. T1 plants were plated on ½ MS containing selection media and selected at 11DAG.

### Photography

To obtain close-up inflorescence images, a Zeiss Stemi 2000-C stereo microscope with a Zeiss Axiocam 105-color digital camera was used. Images were acquired in Zeiss Zen software and brightness contrasted adjusted in Fiji.

### Microscopy

Live *Arabidopsis* SAMs grown at 17-18°C were imaged shortly after floral transition as previously reported^12,13^. SAMs were inserted into a Petri dish containing 2% agarose and submerged in cold water for dissection. Following dissection, Col-0 and *crn svp* SAMs were inserted into another dish and 20 µL of 1.5 mM propidium iodide (PI) in PIPES buffer (35 mM PIPES, 5 mM EGTA, 2 mM magnesium sulfate, pH = 6.8) was pipetted onto the SAM^110^. SAMs were stained for 1-2 minutes before washing 2x with fresh water. *crn* and *crn svp ft* SAMs were submerged in 1.5 mM PI in PIPES buffer for 10-20 minutes before transferring to a water-containing dish for imaging^13^. A 20x water dipping objective was used to acquire a z-stack for each SAM. SAMs were imaged with a 20x/1.0 NA water-dipping objective on a Zeiss LSM 980 confocal microscope using the Airyscan Multiplex CO-8Y mode.GFP markers were excited with a 488 nm laser and emitted signal collected from 380-548 nm. PI was excited with a 561 nm laser and emitted signal collected from 449-549 nm and 573-627 nm. All images using the same reporters were acquired with identical settings. Following acquisition, each z-stack was processed using AiryScan processing (FastAiryScanSheppardSum CO-8Y: 3.7 (2D, Auto.)) Maximum intensity projections of the AiryScan processed images were generated using Zen Blue for presentation in the figures. Images were cropped to remove excess black space at the edges; in some cases, this space included a pedicel. Brightness settings for GFP channels were set to identical values for all SAM images using the same reporter. Brightness in the PI channel was optimized to each SAM for clearest structural presentation.

### Quantitative real-time PCR Analysis

For qRT-PCR, seedlings were plated on ½ MS, stratified in the dark at 4°C for two days, and grown at 22°C under continuous light until 11DAG. 100 mg tissue from 11DAG seedlings was flash-frozen in liquid nitrogen. Total RNA for three biological replicates was isolated using RNAzol RT according to Sigma specifications and quantified using the RNA ScreenTape assay. cDNA was synthesized from 1 µg RNA using the Protoscript ii First Strand cDNA synthesis kit. An Applied Biosystems QuantStudio 6 Flex System and PowerUp SYBR Green Master Mix was used to perform qRT-PCR. Relative expression of *FT* was calculated using the 2^-ΔΔCT^ method using *PP2A* as the housekeeping gene^103^ (see key resources table for primers.)

### Quantification and Statistical Analysis

Unless otherwise noted, flower formation was quantified by classifying the first 30 flower attempts as normal flowers, terminated flowers, or terminated primordia as described in the Results^12,13^. The first flower attempt was defined as the first flower attempt after most apical axillary meristem before stable flower formation. Occasionally, plants grown at 30°C or with very strong *FT* upregulation in cool temperatures did not produce 30 flowers before senescing; for these individuals, the percentage of normal flowers, terminated flowers, and terminated primordia was calculated relative to the total number of flower attempts for that plant.

As we were primarily interested in specific pairwise comparisons between genotypes, we compared the percent terminated primordia between genotypes using non-parametric Mann-Whitney tests and corrected for multiple comparisons using the Holm-Sidak method, with p < 0.05 for the entire family of comparisons. We chose the more conservative Mann-Whitney test as percentages are not normally distributed but were necessary to use to compare data from plants that produced less than 30 flowers. The adjusted p value was used to report statistical significance using asterisks (p < 0.05 * ; p < 0.01 ** ; p < 0.001 *** ; p < 0.0001 ****.) The percent terminated primordia were analyzed using GraphPad Prism v10.4.1. At least two biological replicates were quantified for all experiments. Representative replicates are shown in the figures with sample sizes indicated in the figure legends.

## Acknowledgments

The authors would like to thank George Coupland, Markus Schmid, Tai-ping Sun, Doris Wagner, Marcel Proveniers, Hao Yu, and Benjamin F Holt III for sharing seeds, vectors, and reporters. The Nimchuk lab would like to thank Jamie Winshell for assistance with lab management, James Garzoni and UNC greenhouse staff for assistance with plant growth conditions, Nathanaël Prunet and the Biology Microscopy Core for assistance with imaging, and Corbin Jones for discussions on statistical analysis. The authors wish to acknowledge FLOR-ID for helpful summaries of all flowering time pathways and related research^111^. Work in the lab of Z.L.N was supported by grants from the US National Institutes of Health (R35GM119614) and the National Science Foundation (IOS-1546837). E.S.S was supported by an NSF GRFP fellowship (DGE-2040435). V.C.G. was supported by an EMBO long term fellowship (ALTF 293-2013). Work in the lab of C.F. was supported by a grant from the Swiss National Science Foundation (310030_200318). Fig 7 *Arabidopsis* cartoon created in BioRender. Smith, E. (2025) https://BioRender.com/l67l924

## Author Contributions

E.S.S. conceptualized and performed experiments, analyzed data, acquired support funding, and wrote the manuscript. A.J. and D.S.J performed initial experiments. A.C.W assisted in imaging.

V.C.G. and C.F. provided seed resources. Z.L.N. conceptualized the project and experiments, analyzed data, acquired funding, and wrote the manuscript.

---

**Figure S1:**
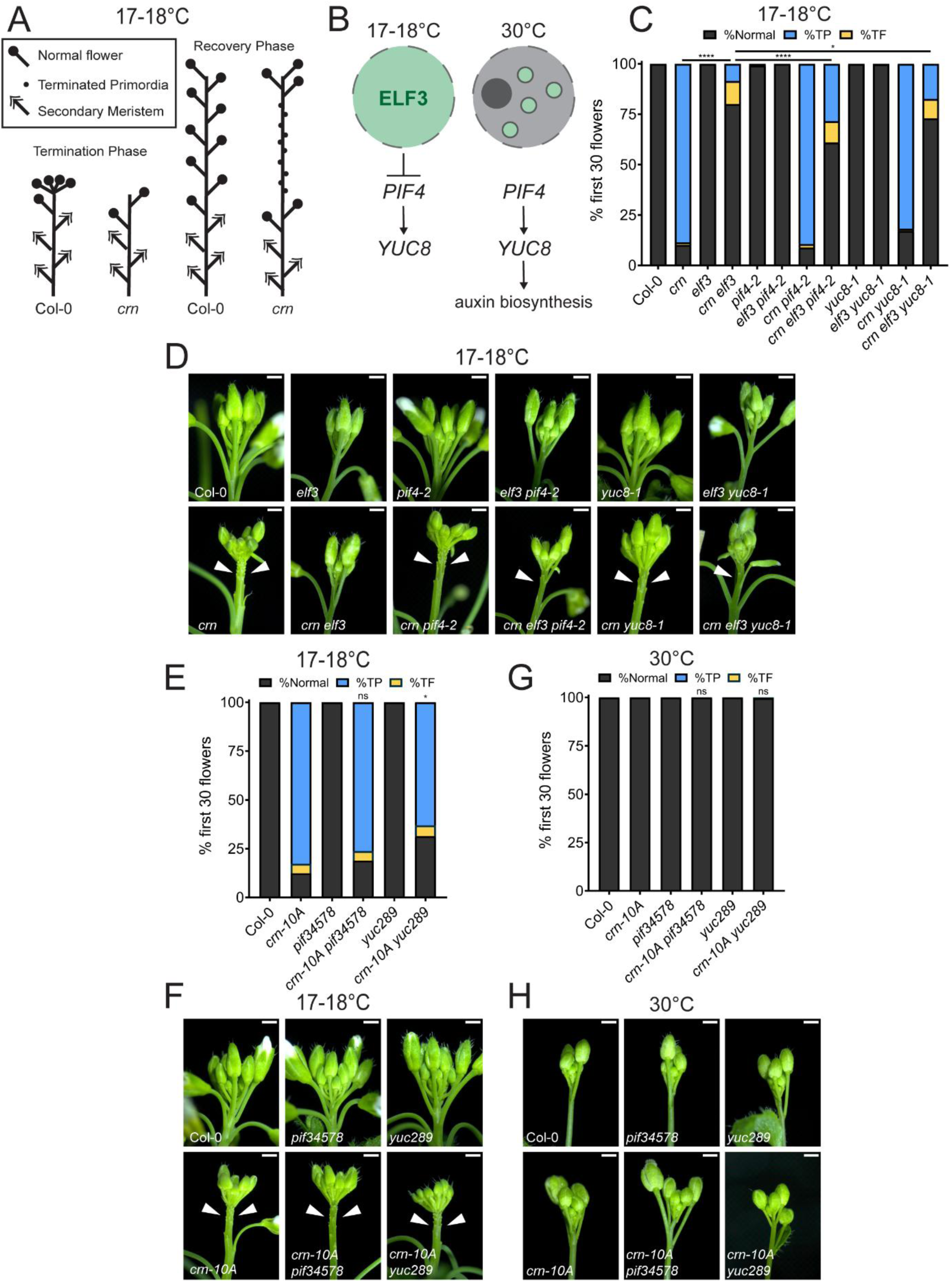
The transcriptional circuitry required for heat-induced auxin biosynthesis is dispensable for canalized flower formation at elevated temperatures. Related to Figure 1. (A) Cartoon of Col-0 and *crn* plants at the termination phase and recovery phases of inflorescence maturation at cooler temperatures. (B) Schematic showing ELF3 protein (Green) nuclear distribution and effects on downstream transcriptional circuit at cooler (17-18°C) and elevated (30°C) temperatures. (C) Quantification of terminated floral primordia by characterizing the first 30 organ attempts as normal (black), terminated primordia (blue), or terminated flower (yellow). Genotypes examined were Col-0 (n=18), *crn* (n=18), *elf3* (n=18), *crn elf3* (n=16), *pif4-2* (n=16), *elf3 pif4-2* (n=14), *crn pif4-2* (n=18), *crn elf3 pif4-2* (n=17), *yuc8-1* (n=16), *elf3 yuc8-1* (n=18), *crn yuc8-1* (n=15), and *crn elf3 yuc8-1* (n=18) grown at cooler temperatures (17-18°C). Statistical significance between pairwise comparisons are indicated on the graph. (D) Representative inflorescence images of all genotypes quantified in (C). (E) Quantification and (F) representative inflorescence images of Col-0 (n=17), *crn-10A* (n=17)*, pif34578* (n=17)*, crn-10A pif34578* (n=17)*, yuc289* (n=18), and *crn-10A yuc289* (n=18) grown at 17-18°C. (G) Quantification and (H) representative inflorescence images of floral primordia termination in Col-0 (n=17), *crn-10A* (n=16)*, pif34578* (n=18)*, crn-10A pif34578* (n=12)*, yuc289* (n=13), and *crn-10A yuc289* (n=12) grown at 30°C. For (E) and (G), statistical significance of *crn-10A* compared to *crn-10A pif34578* or *crn-10A yuc289* is indicated on the graph. Statistical significance between %terminated primordia in indicated pairwise comparisons was calculated with multiple Mann-Whitney tests using the Holm-Šídák method for correcting multiple comparisons. Significance is represented on the graph using asterisks (p < 0.05 *; p < 0.01 **; p < 0.001 ***; p < 0.0001****.) In (D), (F), and (H), terminated primordia are indicated with arrowheads and all scale bars are 1mm.

**Figure S2:**
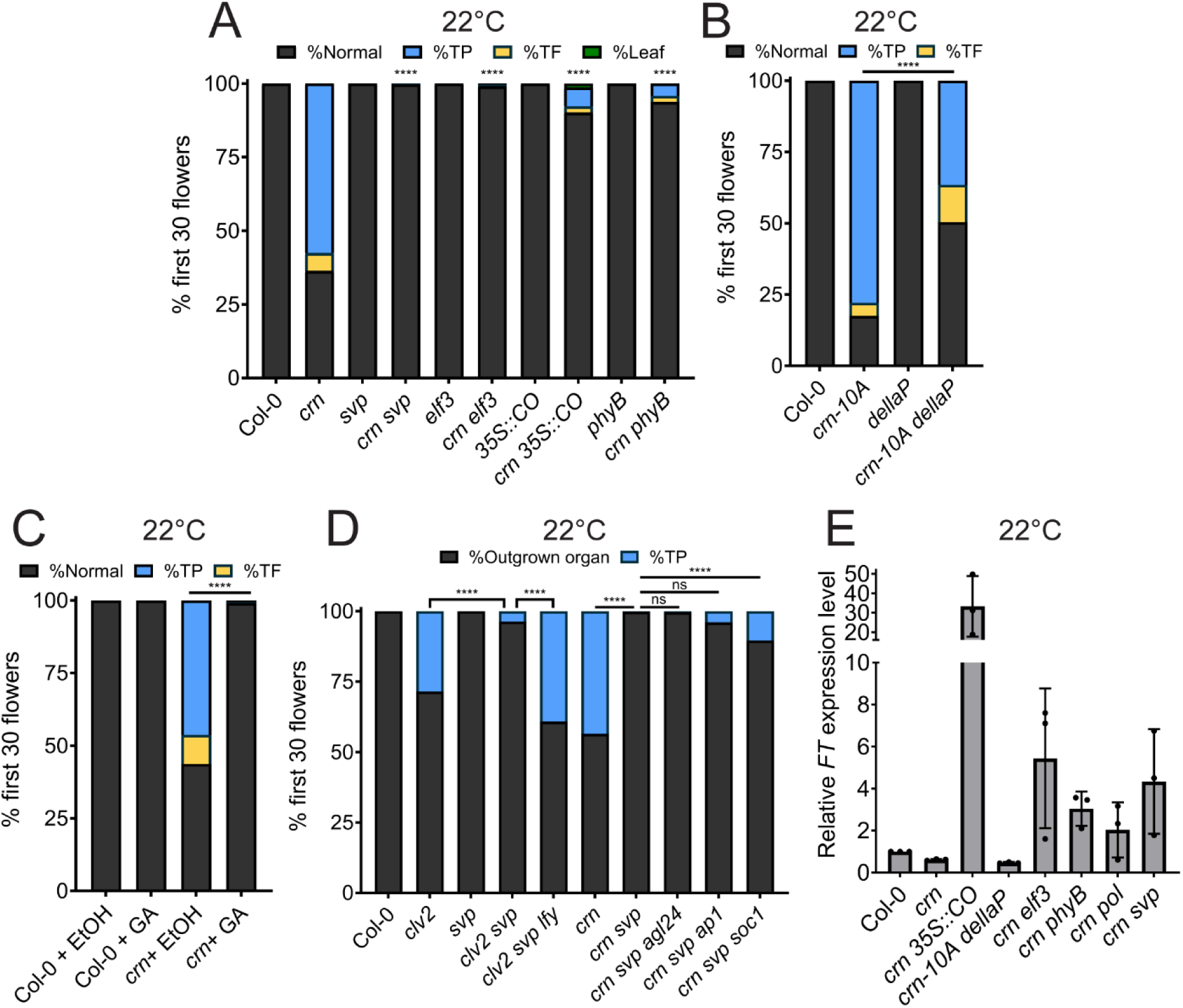
Acceleration of the floral transition restores primordia formation to *crn* mutants at ambient (22°C) temperatures. Related to Figure 2. (A) Quantification of floral primordia termination in Col-0 (n=17), *crn* (n=12), *svp* (n=13), *crn svp* (n=17), *elf3* (n=18), *crn elf3* (n=17), *35S::CO* (n=17), *crn 35S::CO* (n=13), *phyB* (n=18), and *crn phyB* (n=16) grown at ambient temperatures (22°C). (B) Quantification of floral primordia termination in Col-0 (n=16), *crn-10A* (n=16), *dellaP* (n=18), and *crn-10A dellaP* (n=14) plants grown at 22°C. (C) Quantification of floral primordia termination in Col-0 and *crn* plants germinated on 10µM GA_4_ or ethanol (EtOH) containing plates for 11DAG and grown at 22°C. Sample sizes are Col-0 + EtOH (n=16), Col-0 + GA (n=15), *crn* + EtOH (n=18), and *crn* + GA (n=15.) (D) Quantification of floral primordia termination in Col-0 (n=18), *clv2* (n=18), *svp* (n=18), *clv2 svp* (n=18), *clv2 svp lfy* (n=18), *crn* (n=18), *crn svp* (n=16), *crn svp agl24* (n=18), *crn svp ap1* (n=16), and *crn svp soc1* (n=17) plants grown at 22°C. All organs that were not terminated primordia (normal flowers, flower with inflorescence reversion, flowers with identity defects, leaves, and terminated flowers) were classified as “outgrown organs.” Statistical significance between % terminated primordia in indicated genotypes were calculated for (A-D) with multiple Mann-Whitney tests using the Holm-Šídák method for correcting multiple comparisons. Significance is represented on the graph using asterisks (p < 0.05 *; p < 0.01 **; p < 0.001 ***; p < 0.0001****.) (E) Quantitative real-time PCR of relative *FT* expression in 11DAG seedlings of *crn, crn 35S::CO, crn-10A dellaP, crn elf3, crn phyB, crn pol,* and *crn svp* compared to Col-0 seedlings grown at 22°C in continuous light. Three biological replicates are shown. Error bars represent standard deviation.

**Figure S3:**
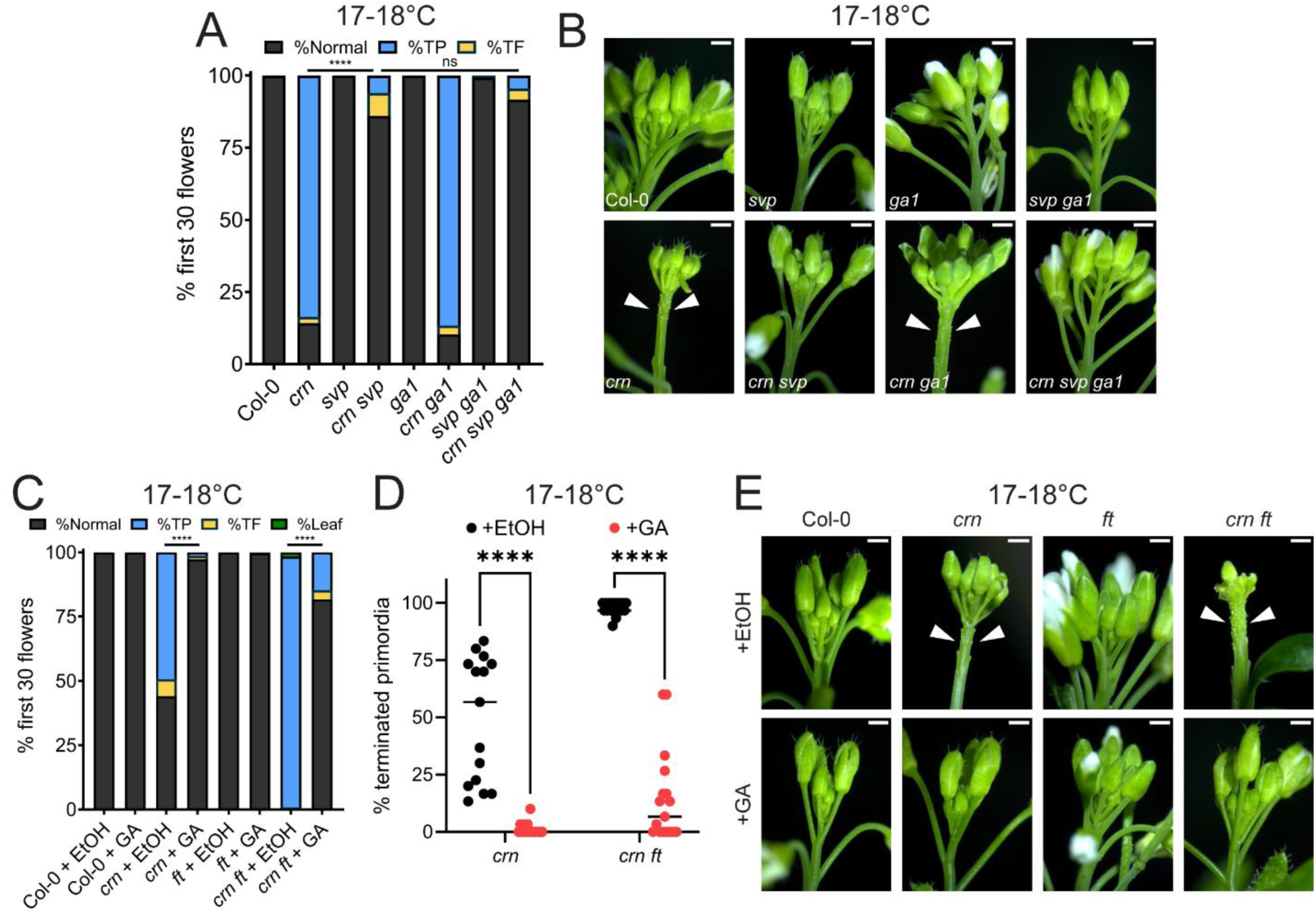
Elevated GA and *FT* restore primordia formation to *crn* in independent pathways. Related to Figure 3. (A) Quantification of floral primordia termination after germination on 10µM GA_4_ or ethanol (EtOH) containing plates for 11DAG and sprayed twice weekly with 10µM GA_4_ or ethanol solution until flowering. Plants were grown at 17-18°C after transferring to soil. Sample sizes were Col-0 + EtOH (n=18), Col-0 + GA (n=17), *crn* + EtOH (n=15), *crn* + GA (n=15), *ft* + EtOH (n=17), *ft* + GA (n=18), *crn ft* + EtOH (n=17), and *crn ft* + GA (n=17). (B) %Terminated primordia for each of the *crn* + EtOH, *crn* + GA, *crn ft* + EtOH, and *crn ft* + GA individuals summarized in (A). Line represents the median. (C) Representative inflorescence images of genotypes quantified in (A). (D) Quantification and (E) representative inflorescence images of terminated floral primordia in Col-0 (n=17), *crn* (n=18), *svp* (n=18), *crn svp* (n=16), *ga1* (n=16), *crn ga1* (n=18), *svp ga1* (n=15), *crn svp ga1* (n=12). Statistical significance between %terminated primordia in indicated genotypes were calculated for (A) and (D) with multiple Mann-Whitney tests using the Holm-Šídák method for correcting multiple comparisons. Significance is represented on the graph using asterisks (p < 0.05 *; p < 0.01 **; p < 0.001 ***; p < 0.0001****.) Arrowheads in (C) and (E) indicate terminated floral primordia. All scale bars are 1mm.

**Figure S4.**
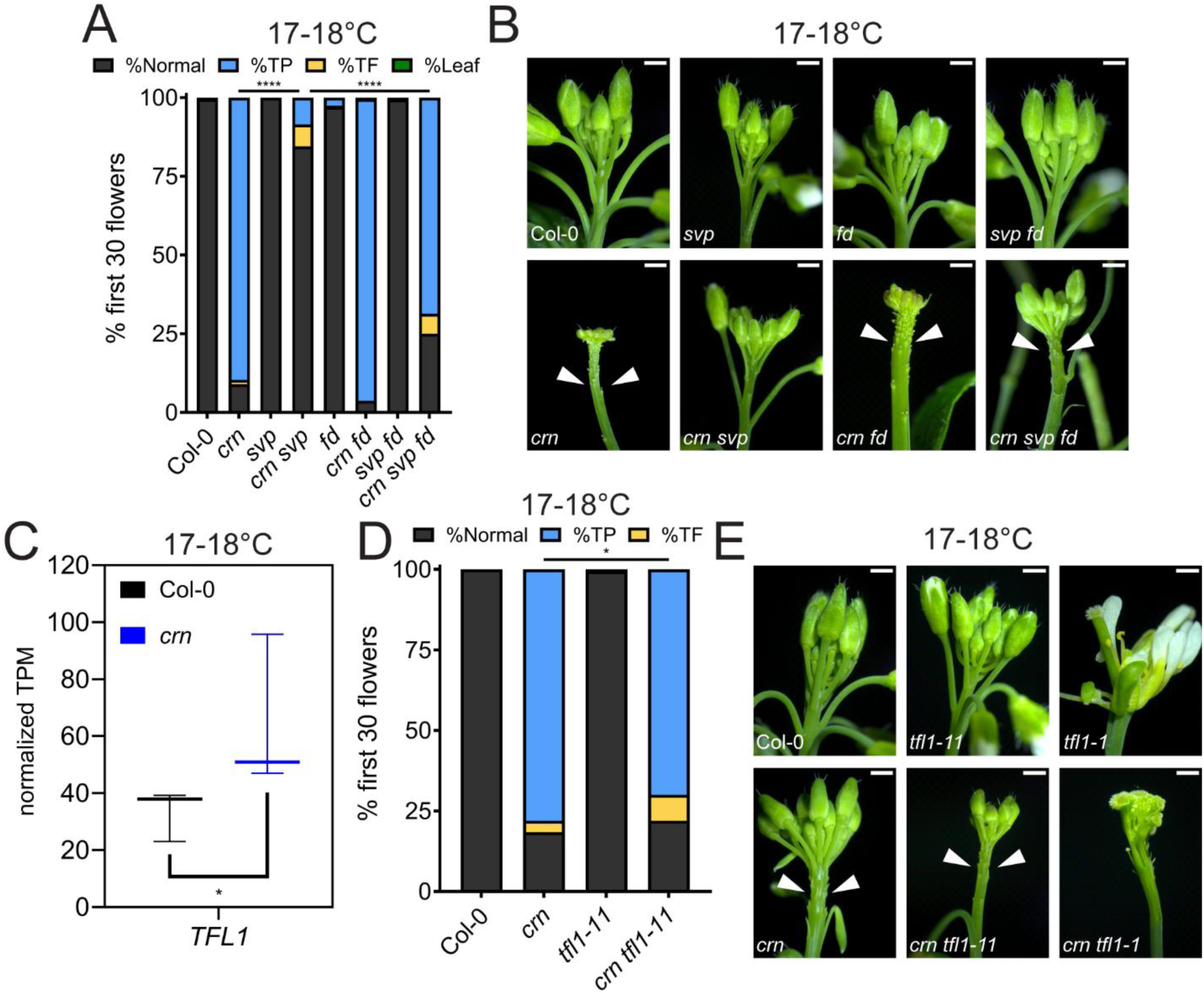
*FD* is required for *FT-*dependent restoration of primordia outgrowth in *crn* mutants. Related to Figure 4. (A) Quantification and (B) representative inflorescence images of floral primordia termination in Col-0 (n=16), *crn* (n=17), *svp* (n=16), *crn svp* (n=9), *fd* (n=18), *crn fd* (n=16), *svp fd* (n=12) and *crn svp fd* (n=18) plants grown at 17-18°C. (C) Expression levels of *TFL1* from RNA-Seq analysis in Col-0 and *crn* SAMs grown at 17-18°C. Normalized transcripts per million (TPM) summed for all the only *TFL1* isoform for three biological replicates are plotted from Kallisto/Sleuth RNA-Seq analysis (p=0.031949). The line indicates the median. (D) Quantification and (E) representative inflorescence images of floral primordia termination in Col-0 (n=17), *crn* (n=17), *crn tfl1-11* (n=16), and *crn tfl1-11* (n=17) plants grown at 17-18°C. *tfl1-1* and *crn tfl1-1* were imaged but not quantified due to the nature of the terminal flower allele obscuring primordia termination. Statistical comparison is indicated on the graph. Statistical significance between %terminated primordia in indicated genotypes were calculated for (A and D) with multiple Mann-Whitney tests using the Holm-Šídák method for correcting multiple comparisons. Significance is represented on the graph using asterisks (p < 0.05 *; p < 0.01 **; p < 0.001 ***; p < 0.0001****.) Arrowheads in (B) and (E) indicate terminated floral primordia. All scale bars are 1 mm.

**Figure S5.**
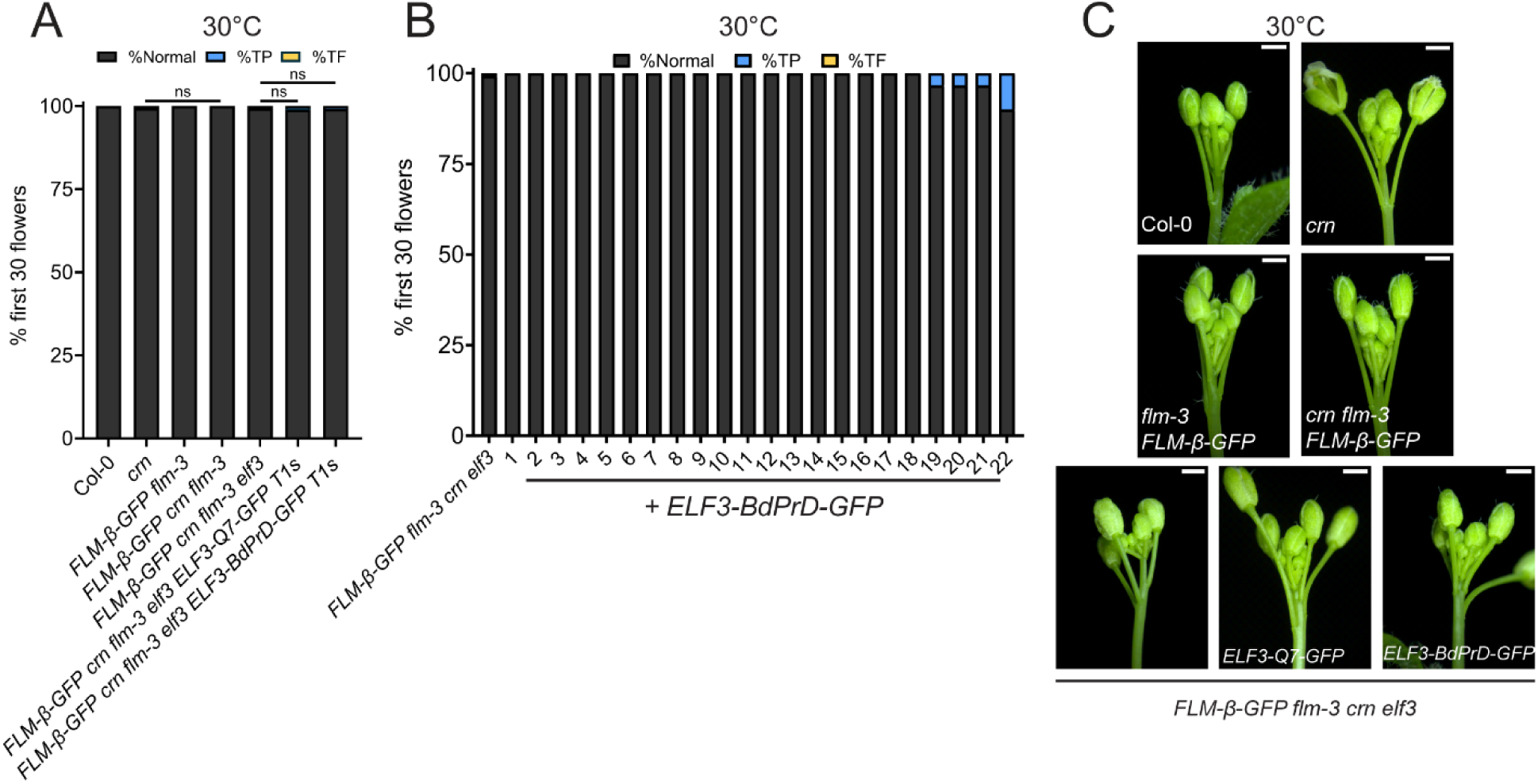
Combined ELF3 and SVP thermosensing is not sufficient to explain heat-mediated primordia restoration in *crn* plants. Related to Figure 5. (A) Quantification of floral primordia termination in Col-0 (n=19), *crn* (n=15), *FLM-β-GFP flm-3* (n=19), *FLM-β-GFP crn flm-3* (n=15), *FLM-β-GFP crn flm-3 elf3* (n=17), *FLM-β-GFP crn flm-3 elf3 ELF3-Q7-GFP* T1s (n=23), and *FLM-β-GFP crn flm-3 elf3 ELF3-BdPrD-GFP* T1s grown at 30°C. (B) Floral primordia termination in *FLM-β-GFP crn flm-3 elf3 ELF3-BdPrD-GFP* T1s summarized in (A). (C) Representative inflorescence images of individuals of genotypes quantified in (A). Statistical significance between %terminated primordia in indicated genotypes were calculated for (A) with multiple Mann-Whitney tests using the Holm-Šídák method for correcting multiple comparisons. Significance is represented on the graph using asterisks (p < 0.05 *; p < 0.01 **; p < 0.001 ***; p < 0.0001****.) All scale bars are 1mm.

